# Meta-PseU: A Meta-Classifier for Robust Prediction of RNA Pseudouridine Modification Sites from Long Sequences

**DOI:** 10.64898/2025.12.08.693080

**Authors:** Takumi Suto, Md. HARUN-OR-ROSHID, Hiroyuki Kurata

**Author notes:** Corresponding authors: Hiroyuki Kurata.

## Abstract

Pseudouridine (Ψ) represents one of the most abundant and evolutionarily conserved RNA modifications. Ψ provides an additional hydrogen-bond donor that enhances RNA structural stability and modulates translation. It participates in diverse biological processes, including RNA-protein interactions, splicing, translational control, and stress responses. Aberrant pseudouridylation is implicated in cancer, neurodegenerative disorders, and autoimmune diseases. Despite its biological importance, experimental identification of Ψ sites remains time-consuming and costly, limiting the feasibility of transcriptome-wide profiling. Computational approaches have therefore become essential complements to experimental techniques. However, current machine-learning and deep-learning predictors suffer from limitations such as small datasets and limited generalizability. To overcome these issues, we have constructed new long-sequence datasets derived from RMBase 3.0 and developed Meta-PseU, a logistic regression-based meta-classifier that stacks multiple single-feature or baseline classifiers across three species of human, mouse, and yeast. Meta-PseU substantially reduced the performance gaps between training and independent test datasets, presenting superior generalization. Meta-PseU substantially outperformed state-of-the-art predictors and achieved increasing accuracy with increasing sequence length. This work offers a new framework for robust Ψ-site identification by using long sequences. The datasets and programs are freely accessible at https://github.com/kuratahiroyuki/MetaPseU.

## 1. Introduction

Post-transcriptional chemical modifications of ribonucleic acids (RNAs) play essential roles in regulating gene expression and cellular processes across all domains of life, including eukaryotes, bacteria, and archaea [1, 2]. To date, more than 300 types of RNA modifications have been identified [3, 4]. Among them, pseudouridine (Ψ) is one of the most abundant and evolutionarily conserved modifications [5]. Ψ is an isomer of uridine in which the C–N glycosidic bond is replaced by a C–C bond, providing an additional hydrogen-bond donor that stabilizes RNA structures and contributes to translational regulation [6, 7]. It is widely distributed in various RNA species including mRNA, tRNA, rRNA, snRNA, snoRNA, miRNA, and lncRNA and is involved in various biological processes such as RNA and protein interactions, splicing regulation, translation regulation, and stress responses [8]. Abnormal pseudouridylation can cause many human diseases including cancer, neurodegenerative disorders, and autoimmune disorders [9–11].

Experimental technologies have been developed to identify Ψ sites, such as Pseudo-seq [12], CeU-seq [13], Ψ-seq [14], and RBS-seq [15]. Although these methods based on complex chemical reactions provide nucleotide-level resolution, they are both time-consuming and costly, making them unsuitable for high-throughput, transcriptome-wide analysis. Therefore, computational approaches have become indispensable for complementing experimental identification.

Numerous machine-learning (ML) and deep-learning (DL)-based predictors have been proposed for Ψ-site identification. iRNA-PseU [16], PseUI [17], iPseU-NCP [18], RF-PseU [19], and XG-PseU [20] implemented a range of machine-learning methods combined with diverse encoding schemes, including composition-based, position-order, and physicochemical property representations. EnsemPseU [21] and PseU-ST [22] were designed as an ensemble framework that stacked multiple machine-learning models, each trained on different feature-encoding methods, to improve predictive robustness and overall performance. PseU-FKeERF [23] employed a fuzzy kernel evidence-based Random Forest (RF) classifier, enabling it to capture nonlinear feature interactions while maintaining interpretability. PseUdeep [24] and RSCNN-PseU [25] implemented deep-learning approaches such as convolutional neural networks (CNNs) to automatically learn hierarchical feature representations from sequence data. Most of these models were trained on the common benchmark datasets derived from RMBase 2.0 [26], which include H_990 (human), M_944 (mouse), and S_628 (yeast) for training and H_200 and S_200 for independent testing. Such a shared dataset facilitates direct performance comparisons across models, while the limited sample size poses challenges in assessing generalization capability.

Many classifiers achieved exceptionally high AUC values (e.g., >0.95) on training datasets, but provided poor reproducibility and low generalizability when evaluated on independent test dataset as shown in **Table S1**, where we reproduced and evaluated several state-of-the-art (SOTA) models with publicly available models including PseU-ST [22], PseU-FKeERF [23], and RSCNN-PseU [25]. These SOTAs exhibited high prediction performances on the training datasets of human and yeast, respectively, but provided poor reproducibility and less generalizability on the test datasets. Thus, the reproducibility and generalizability of these models remain to be fully clarified (**Table S2**).

Recently, RMBase 3.0 [3] has been released as a comprehensive resource for decoding the landscape, mechanisms, and functional implications of RNA modifications. This updated database includes a substantially larger number of annotated Ψ sites across various species, which presents an excellent opportunity to refine and extend existing datasets, thereby enabling more robust identification of RNA modification sites.

To solve the above generalizability and limited datasets, we have constructed new long-sequence datasets based on RMBase 3.0 and have developed a novel computational Ψ site predictor named Meta-PseU that integrated multiple single-feature-based models via logistic regression (LR), as shown in **Figure 1**. Meta-PseU considered 17 different feature-encoding schemes combined with 7 ML classifiers and two encoding methods with three DL architectures and substantially outperformed the SOTAs. Meta-PseU extended the sequence length up to 201 instead of a conventional length of 21 to further enhance the prediction performance.

**Figure 1.**
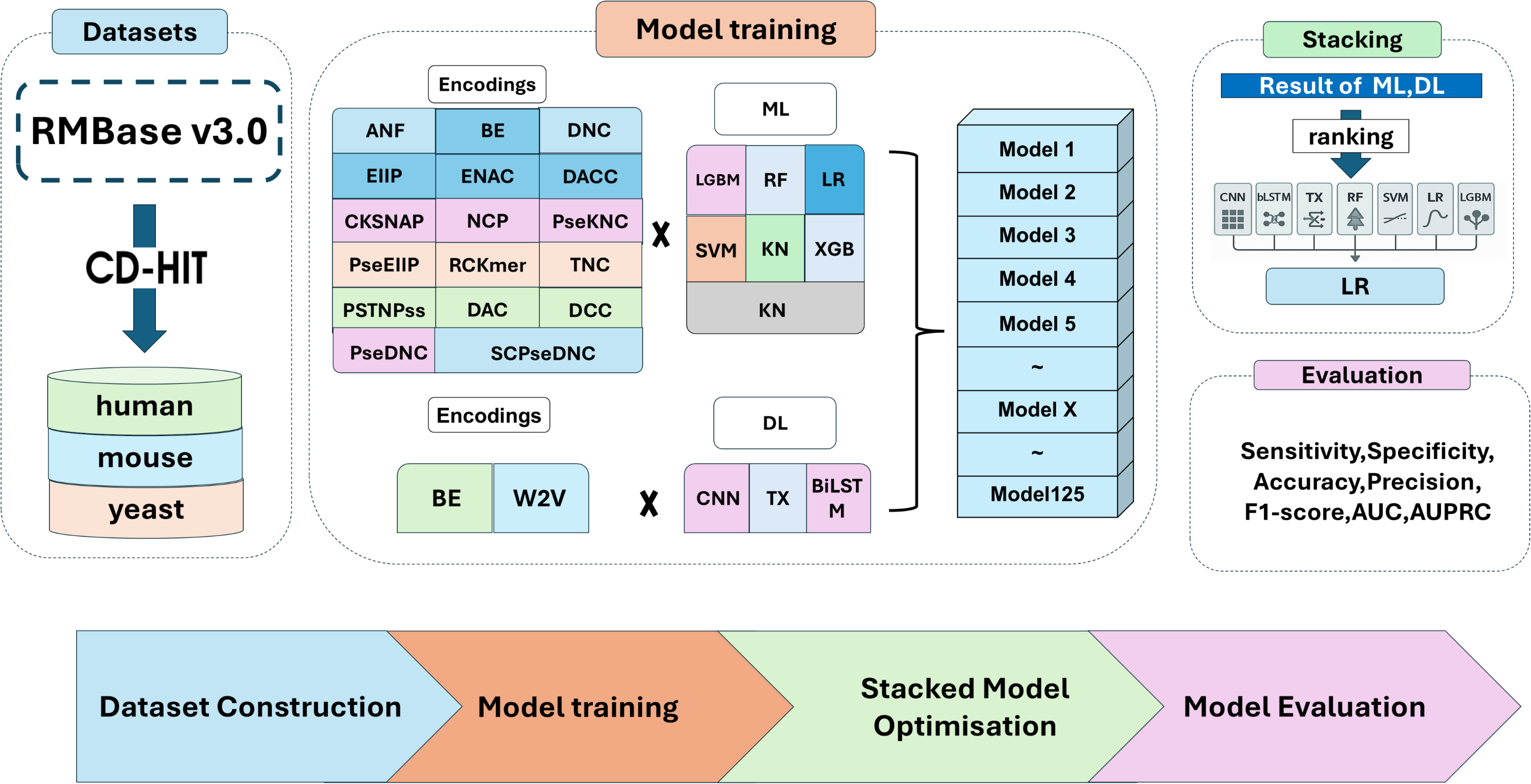
Workflow of Meta-PseU development. The development process comprises four major stages. (i) Construction of benchmark Ψ-site datasets from RMBase 3.0, while removing redundant sequences using CD-HIT. (ii) Construction of baseline models: 17 single encoding method employing-ML models and W2V and OH-based DL architectures. (iii) Construction of meta-classifier stacking the baseline models using logistic regression. (iv) Evaluation of model performance.

## 2. Materials and Methods

### 2.1 Dataset construction

We constructed benchmark datasets for developing and evaluating predictive models of Ψ modification sites in RNA sequences of *Homo sapiens* (human), *Mus musculu*s (mouse), and *Saccharomyces cerevisiae* (yeast). Each dataset was derived from the RMBase v3.0 database [3], which integrates experimentally validated RNA modification sites identified through various high-throughput techniques. For each species, experimentally confirmed Ψ sites were designated as positive samples, whereas non-modified uridine (U) sites within the same genes were randomly selected as negative samples. Each modification site was mapped to the reference genome of the respective species, and the corresponding RNA sequence was extended to 201 nucleotides in length, with the Ψ site positioned at the center. From these 201-nucleotide sequences, the datasets with different sequence lengths of 41, 81, 121, 161, and 201 were constructed by trimming both ends, enabling a systematic evaluation of how sequence length influences prediction performance. To eliminate sequence redundancy, sequences were clustered using CD-HIT [27] with a 70% sequence identity threshold, and only non-redundant sequences were retained. To prevent class imbalance during model training, a ratio of positive to negative samples was adjusted to 1:1. Finally, each dataset was divided into training (80%) and independent test (20%) sets, and this partitioning was consistently applied across all models developed in this study. A statistical summary of the constructed datasets is shown in **Table 1**.

**Table 1.**
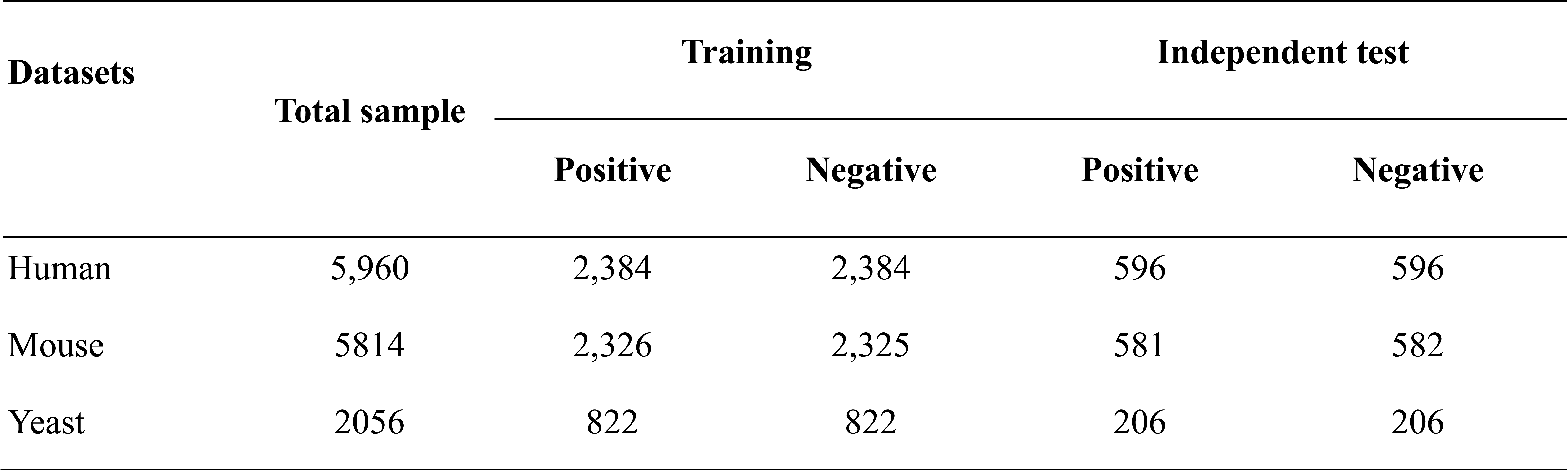
Datasets of Ψ site modification in human, mouse and yeast.

### 2.2. Encoding methods

Feature encoding methods play a crucial role in constructing sequence-based classifier [28, 29].We employed 17 different sequence-encoding methods which are roughly grouped into the composition-based encodings, position-order-based encodings, position- and composition-based encodings, correlation-based encodings, and language models.

#### Composition-based encodings

##### Nucleic Acid Composition (NAC)

The NAC encoding calculates the frequency of each nucleotide type (A, C, G, U) in an RNA sequence[30]. It simply counts the number of occurrences of each nucleotide and divides it by the total sequence length. The frequency of nucleotide is defined as:

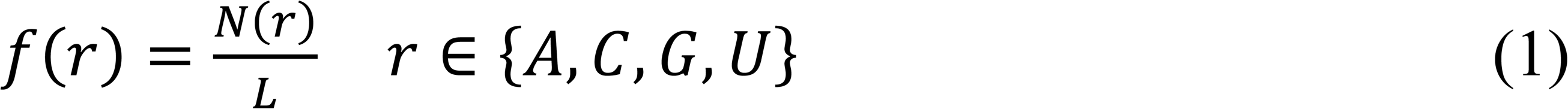

where *N*(*r*) represents the number of occurrences of nucleotide *r* and *L* is the total sequence length. This NAC provides a global statistical representation of nucleotide distribution within an RNA sequence.

##### Di-Nucleotide Composition (DNC)

The DNC extends NAC by considering pairs of adjacent nucleotides (dinucleotides). It computes the frequency of each possible dinucleotide combination within the RNA sequence, which captures local dependencies between neighboring bases. The DNC feature is defined as:

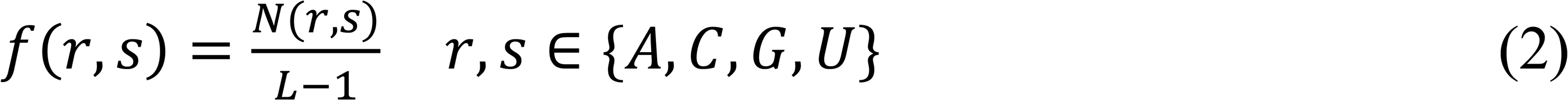

where *N*(*r*, *s*) is the number of occurrences of dinucleotide (*r*, *s*) in the sequence. There are 16 possible dinucleotide combinations in total (4² = 16).

##### Tri-Nucleotide Composition (TNC)

The TNC encoding further extends NAC by considering triplets of adjacent nucleotides (trinucleotides). It captures higher-order local dependencies across three consecutive bases. The TNC frequency is computed as:

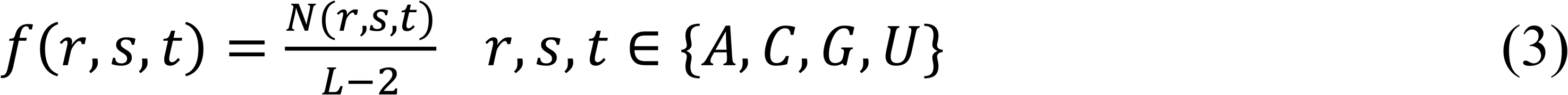

where *N*(*r*, *s*, *t*) denotes the number of occurrences of trinucleotides (*r*, *s*, *t*). There are 64 possible trinucleotide combinations (4³ = 64).

##### Composition of K-spaced Nucleotide Pairs (CKSNAP)

The CKSNAP encoding is an extension of the DNC method [31]. It calculates the frequency of nucleotide pairs separated by an interval of K nucleotides. Each pair is counted and divided by the total number of possible pairs, providing a normalized representation. CKSNAP captures correlations between non-adjacent nucleotides and reflects the spatial dependency between bases, which can be important for recognizing modification-related motifs.

##### RCKmer (Reverse Complement K-mer)

The RCKmer encoding is a variant of the traditional K-mer approach that considers reverse complement symmetry in nucleotide sequences [31]. In standard K-mer encoding, each K-length substring is treated as a unique feature, which can lead to redundancy because the reverse complement of one K-mer often conveys equivalent biological information. In the RCKmer scheme, each K-mer and its reverse complement are merged into a single feature, reducing the dimensionality and improving biological interpretability. For RNA sequences, there are 16 possible dinucleotides (2-mers): AA, AC, AG, AU, CA, CC, CG, CU, GA, GC, GG, GU, UA, UC, UG, UU. By collapsing reverse complements (e.g., “AA” ↔ “UU”, “AGU” ↔ “ACU”, “GUCAU” ↔ “AUGAC”), the RCKmer representation for dinucleotides results in 10 distinct features: AA, AC, AG, AU, CA, CC, CG, GA, GC, and UA. The RCKmer captures both forward and reverse sequence composition patterns while avoiding redundancy caused by complementary strand equivalence.

##### Pseudo Electron–Ion Interaction Potential (PseEIIP)

The PseEIIP encoding extends the basic EIIP representation by incorporating trinucleotide composition into the feature space [32]. Each nucleotide (A, C, G, U) is assigned to its corresponding EIIP value (A = 0.1260, C = 0.1340, G = 0.0806, U = 0.1335), and the average EIIP value of each trinucleotide is calculated as follows:

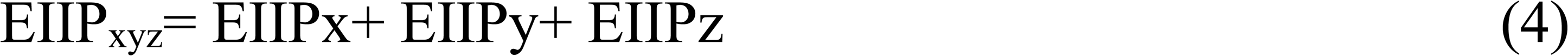

The normalized frequency of each trinucleotide, f_xyz_ is then multiplied by its corresponding EIIP_xyz_ value to form the final feature vector:

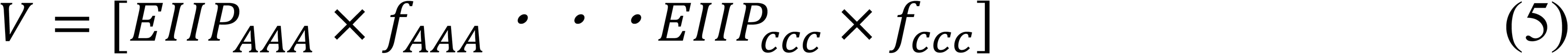

This representation integrates both electronic properties and trinucleotide frequency, allowing the model to capture local sequence structure together with the physical and chemical context underlying Watson–Crick base pairing interactions.

#### Position-based encodings

##### One-hot (OH)

The OH represents each nucleotide as a four-dimensional binary vector. Specifically, A, U, G, and C are encoded as (1, 0, 0, 0), (0, 1, 0, 0), (0, 0, 1, 0), and (0, 0, 0, 1), respectively. Thus, an RNA sequence of *L* nucleotides can be represented as a 4 × *L*–dimensional feature vector, where each position preserves the categorical identity of the nucleotide. The OH is simple yet effective for representing the discrete nature of RNA sequences.

##### Nucleotide Chemical Property (NCP)

The NCP encoding represents each nucleotide (A, C, G, U) based on its fundamental chemical and structural characteristics [31]. Since nucleotides can be classified into three physicochemical groups, each nucleotide can be represented as a 3-dimensional vector: A = (1, 1, 1), C = (0, 1, 0), G = (1, 0, 0), U = (0, 0, 1). This encoding allows chemical diversity to be numerically represented, capturing both the physical and bonding characteristics of each base.

##### Electron-Ion Interaction Pseudopotential (EIIP)

The EIIP encoding assigns a specific numerical value to each nucleotide according to its electron–ion interaction potential [32]. Each RNA sequence is then converted into a vector of these values, preserving the biophysical properties of the bases. This physicochemical representation allows models to learn patterns based on the electronic characteristics of nucleotides, which are linked to biological functionality.

#### Position- and composition-based encodings

##### Enhanced Nucleic Acid Composition (ENAC)

The ENAC encoding calculates the local nucleotide composition within a fixed-length sliding window along the RNA sequence [33]. A window of predefined size (*k*) slides continuously from the 5′ to the 3′ end of the sequence, and the NAC is computed for each window position. By default, the window size is *k* = 5. This process generates a series of local composition profiles that capture regional variations in nucleotide frequency, allowing the model to learn position-dependent sequence patterns.

##### Accumulated Nucleotide Frequency (ANF)

The ANF encoding represents the cumulative frequency of each nucleotide along the RNA sequence [34]. It is defined as:

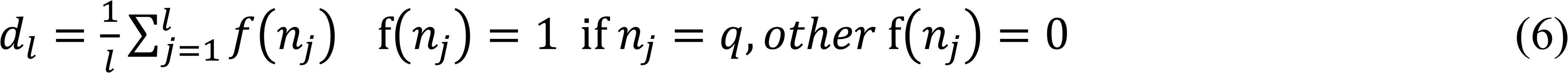

where *d*_*l*_ is the density of the target nucleotide q at position l (1 ≤ l ≤ L). For instance, in the sequence “CACAGUCG”, the density of cytosine (C) at position l = 3 is given by:

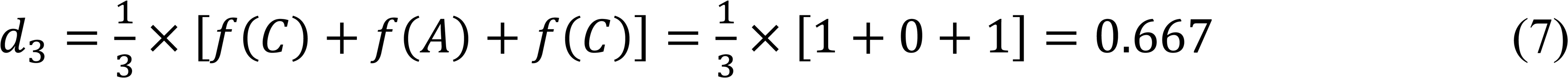

ANF captures the positional accumulation pattern of specific nucleotides.

##### Pseudo K-tuple Nucleotide Composition (PseKNC)

The PseKNC encoding represents an RNA sequence as a numerical feature vector by combining local sequence composition (K-mer frequencies) and global sequence-order information based on physicochemical properties of nucleotides [35]. It starts from K-tuples, e.g., dinucleotides (K=2) or trinucleotides (K=3). Then it adds pseudo components that describe correlations between nucleotides at certain distances using physicochemical properties such as stacking energy and melting temperature. These correlations are controlled by a parameter **λ**, which defines how far apart positions are considered. Briefly, it extends NCP by incorporating both the chemical properties of nucleotides and the long-range sequence-order information.

##### Position-Specific Trinucleotide Propensity based on Single Strand (PSTNPss)

The PSTNPss encoding constructs a position-specific matrix of trinucleotide frequencies for both positive and negative samples [16]. For a sequence of length *L*, the matrix is represented as:

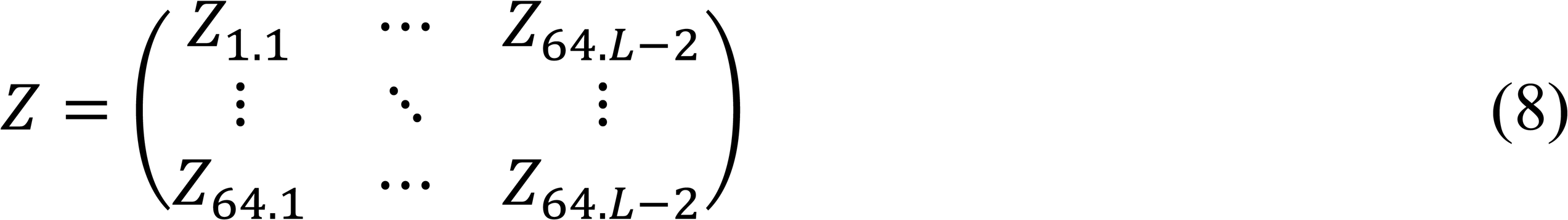

where each element *Z_i,j_* denotes the frequency difference of the *i*th trinucleotide at position j between positive and negative datasets. The feature vector for a given sequence is then constructed as:

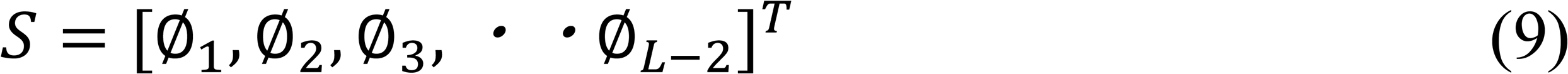

where ∅_*u*_ corresponds to the positional preference score of each trinucleotide. This encoding preserves local sequence context and position-specific preferences for modified versus unmodified sites.

#### Correlation-based encodings

##### Pseudo Di-Nucleotide Composition (PseDNC)

The PseDNC encoding represents RNA sequences by combining DNC with sequence-order correlation factors[36, 37]. Given an RNA sequence *S* = *R*_1_*R*_2_*R*_3_*R*_4_・・・*R*_*L*_

where R_i_ denotes the nucleotide at position i, The basic DNC vector is defined as:

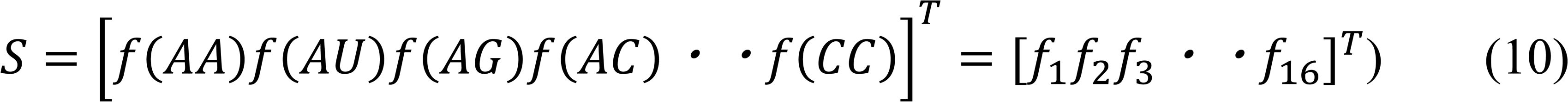

where *f*(*r, s*) represents the normalized frequency of dinucleotide (*r, s*). However, simple composition-based encoding (such as NAC or DNC) loses the sequence-order information. To address this limitation, PseDNC introduces pseudo components that quantify the correlations between di-nucleotide physicochemical properties at different positions, thereby integrating both compositional and sequential dependencies.

##### Parallel Correlation Pseudo Dinucleotide Composition (PCPseDNC)

The PCPseDNC encoding method derives from PseDNC, which considers the correlation between adjacent dinucleotides within an RNA sequence. Unlike PseDNC, PCPseDNC utilizes 38 predefined physicochemical indices instead of the conventional six. It incorporates the parallel correlations between dinucleotides located at positions i and j, as well as correlations among dinucleotides separated by a specific lag *k* [31]. This approach effectively captures both local and global dependencies in nucleotide physicochemical properties.

##### Dinucleotide-Based Cross Covariance (DCC)

The DCC encoding quantifies the correlations between physicochemical indices of two dinucleotides separated by a fixed number of nucleotide (lag) along the RNA sequence [31]. It is defined as:

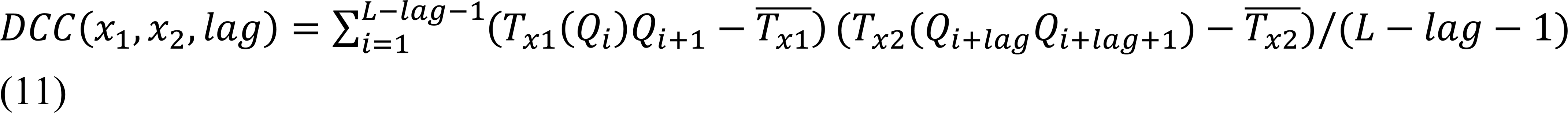

where *T*_x_(Q_i_Q_i+1_) denotes the numerical scale of the physicochemical property *x* for the dinucleotide at position i, 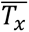 is the mean of that property across the sequence. DCC thus reflects the covariance between local physicochemical features across specified lags, providing insight into how biochemical tendencies propagate along the RNA chain.

##### Dinucleotide-based Auto Covariance (DAC)

The DAC encoding measures the auto correlation between the same physicochemical indices of dinucleotides separated by a fixed distance within a sequence [31]. It is mathematically defined as:

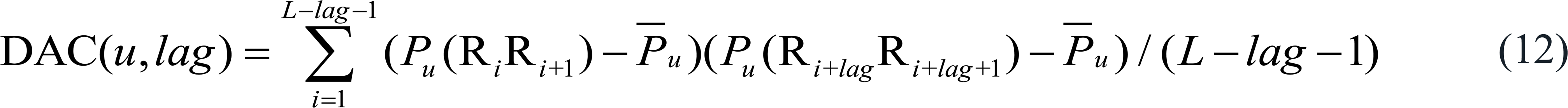

where *x*_1_ *and x*_2_ are distinct physicochemical indices, and *Q*_*i*_*Q*_*i*+1_ represents the dinucleotide at position *i*.

##### Dinucleotide-based Auto-Cross Covariance (DACC)

The DACC encoding is an extension of DAC [34]. Unlike DAC, it effectively captures both the auto-covariance and cross-covariance relationships among biochemical features within RNA sequences, providing a richer representation of their underlying structural and functional patterns.

#### Word2vec (W2V)

W2V is an essential feature embedding method based on natural language processing is extensively employed in text data analysis and has proven particularly effective in various pattern recognition tasks involving sequence data [38, 39]. In this study, we employed the skip-gram algorithm to train a W2V model on the RNA sequences retrieved from RNAcentral [40] using the key words “Human”, “Rfam”, and “non-coding RNA”. This resulted in 128-dimensional feature vectors for each nucleotide.

### 2.3. Learning methods

We implemented seven ML and three DL models to construct a predictive framework for identifying RNA Ψ modification sites. The ML classifiers included four tree-based algorithms: RF, Extreme Gradient Boosting (XGB), and Light Gradient Boosting Machine (LGBM), two decision boundary-based classifiers: Support Vector Machine (SVM) and LR, and two probabilistic or distance-based methods: Naïve Bayes (NB) and K-Nearest Neighbor (KNN) [41]. For the DL models[42], we constructed CNN, Bidirectional Long Short-Term Memory (BiLSTM), and Transformer encoder (TX) architectures [43]. A brief overview of these classifiers [44] is provided below.

RF is one of the most popular supervised ML algorithms, widely used in bioinformatics for various classification and prediction problems[45]. It constructs multiple decision trees trained on different subsets of the training data and predicts the class of new samples by majority voting. SVM is a robust supervised learning algorithm frequently applied to classification problems. It identifies an optimal hyperplane that separates data points into two classes in a high-dimensional space. By applying kernel functions, SVM can handle complex and nonlinear relationships, making it highly effective for high-dimensional and heterogeneous biological data [46]. XGB is a powerful ensemble-based algorithm designed for both regression and classification tasks, particularly effective on large and sparse datasets [47]. It achieves high accuracy by combining multiple decision trees trained via gradient boosting. To mitigate overfitting, XGB employs L1 and L2 regularization to penalize model complexity. Its scalability and parallelized implementation make it especially suitable for computationally intensive bioinformatics analyses. LGBM is an efficient gradient boosting algorithm characterized by fast training speed, low memory usage, and excellent scalability [48]. It integrates several optimization strategies such as sparse feature handling, early stopping, and regularization. The NB classifier is a probabilistic ML model based on Bayes’ theorem, assuming conditional independence among features. The KNN algorithm classifies samples based on the majority class among their K nearest neighbors in the training set, determined using Euclidean distance. The choice of K greatly affects performance; hence, it was optimized through cross-validation. LR is a generalized linear model commonly used for binary classification problems due to its interpretability and robustness [44].

CNNs extend a multilayer perceptron by using locally connected receptive fields to capture spatially correlated features. Through stacked convolution and pooling layers, they learn increasingly abstract representations, making them effective for detecting sequence motifs and structural patterns[43, 49]. LSTM networks address the vanishing-gradient problem in RNNs by using memory cells and gating mechanisms to learn long-range dependencies [50]. BiLSTMs extend LSTMs by processing sequences in both forward and backward directions, enabling the capture of contextual information from both past and future positions. The TX network is a DL architecture based on the Transformer encoder, which uses self-attention instead of recurrent or convolutional operations [51]. This attention-only design enables parallel computation, faster training, and strong generalization. The TX uses three stacked encoder blocks to learn contextual relationships and long-range dependencies among nucleotides. The aggregated outputs from local sub-sequences are passed through a sigmoid function for binary classification

### 2.4. Meta-learning approach

To construct the final integrated predictor, we adopted a meta-learning (stacking) framework. A total of 125 baseline models were generated by combining 17 feature-encoding methods with 7 ML algorithms and 3 DL architectures. Each baseline model produced prediction-probability scores through 5-fold cross-validation on the training dataset[29]. These probability scores were then used as input features for an LR-based meta-predictor, which learned to optimally integrate the outputs of the baseline models. The LR model can be expressed as

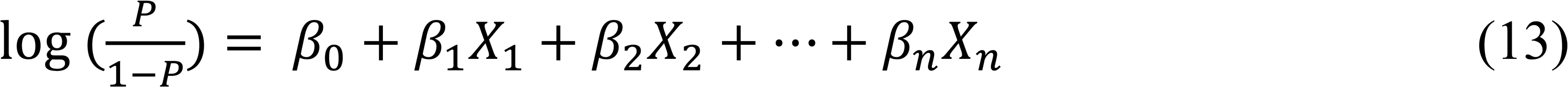

where β_0_ and β_i_ are regression coefficients, *X*_*i*_ is the probability score generated by the *i*th baseline model.

### 2.5. Performance evaluation

To evaluate the performance of the prediction models, eight statistical measures were employed: sensitivity (SEN), specificity (SPE), accuracy (ACC), Matthews correlation coefficient (MCC), and area under the receiver operating characteristic curve (AUC). Most compatible measures were defined as

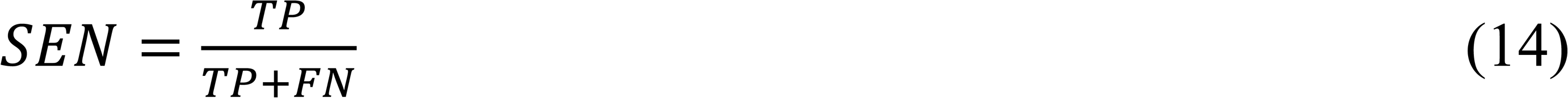

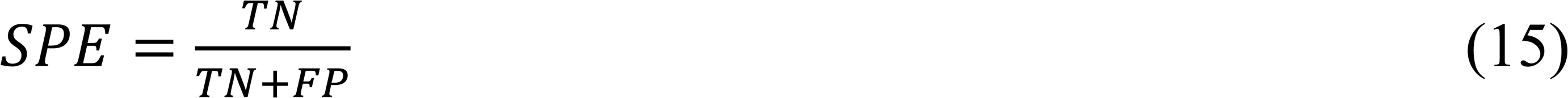

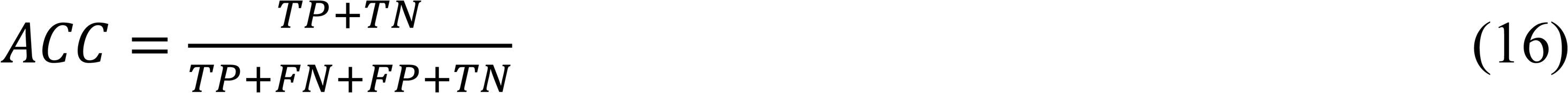

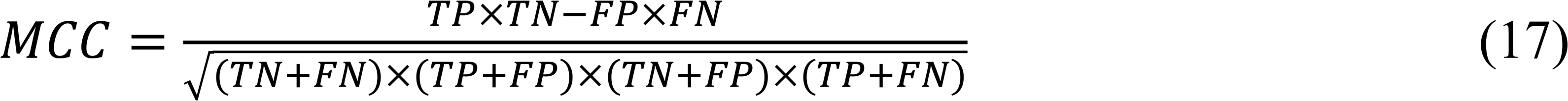

where TP is the number of true positives, FP is the number of false positives, TN is the number of true negatives, and FN is the number of false negatives.

## 3. Results

### 3.1. Nucleotide preference analysis

**Figure 2** illustrates the sequence context surrounding Ψ sites using pLogo analysis. The upper panel represents positive sequences, while the lower panel depicts negative control sequences. In both panels, the upper half of each pLogo shows the nucleotides that are significantly overrepresented, whereas the lower half shows those that are underrepresented relative to a background model. In humans we observed a distinct nucleotide composition bias surrounding Ψ sites in the positive sequences: strong G- and U-enrichment upstream, preferential C and G downstream context, and significant depletion of A and G near the modification site. In mouse, A and U were overrepresented and C and G were underrepresented at position 1, whereas C and U were overrepresented and A and G were underrepresented at position 2 in the positive sequences. In yeast, G, U, G, and G were overrepresented and A, A, C, and T were underrepresented at positions of -3, -2, -1, and 2, respectively. In contrast, the negative sequences for all the three species displayed a lack of clear overrepresentation near the central position. The strong depletion signal of A at position of 1 was observed across three species, suggesting avoidance of A in Ψ-modified contexts, while the other regions presented relatively uniform nucleotide frequencies, suggesting no strong sequence bias in the negative sequences. Nucleotide preference analysis demonstrates that some motif enrichments in the positive sequences are specific to Ψ modification rather than random base occurrence, providing mechanistic insight into enzyme recognition and serving as a basis for improved Ψ site prediction algorithms.

**Figure 2.**
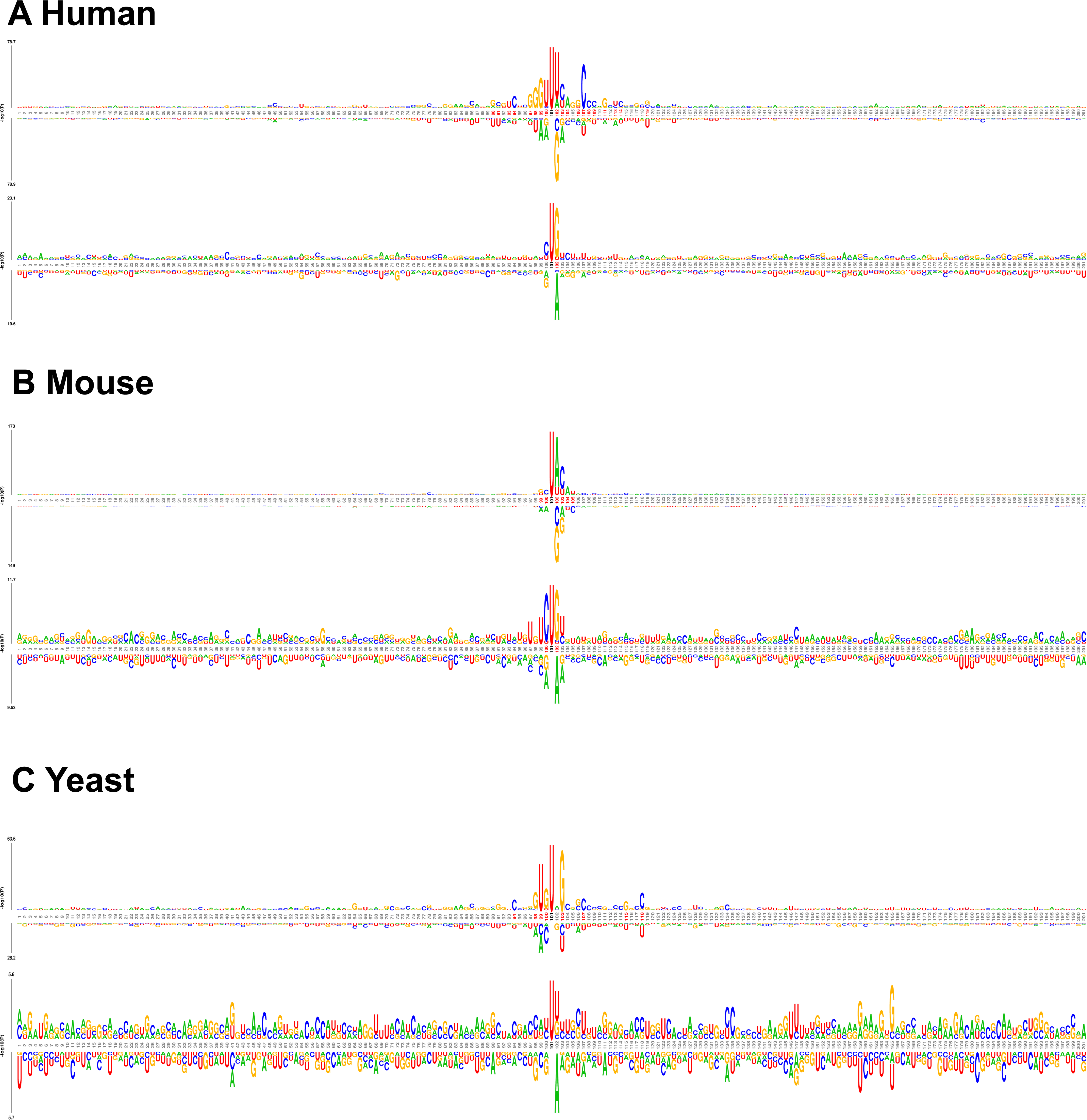
Nucleotide preference pattern analysis around Ψ modification sites by pLogo for human (A), mouse (B), and yeast (C). The upper panel represents positive sequences, while the lower panel depicts negative control sequences. In both panels, the upper half of each pLogo shows nucleotides that are significantly overrepresented, whereas the lower half shows those that are underrepresented relative to a background model

### 3.2. ML performance trends with varying sequence lengths

To evaluate the prediction performance, we analyzed three species datasets of human, mouse, and yeast using seven ML algorithms (LGBM, RF, XGB, SVM, KNN, LR, and NB) implementing 17 different encoding methods. Figure 3 illustrates the changes in AUC with increasing sequence lengths (41, 81, 121, 161, and 201) for all 17 encodings used by LGBM-based classifiers. The prediction performance for the other 6 ML models is provided in **Figures S1-S6**. Across all encoding methods, AUC values were a little higher in the training datasets than in the test datasets, reflecting slight overfitting. OH, NCP, and EIIP consistently achieved very high AUCs (0.8∼0.82) across species, indicating strong discriminative power. The difference in AUCs between training and testing was small (mostly <0.02) for the well-performing encodings, suggesting reasonable generalizability. In contrast, ANF and PSTNPss yielded very poor performance (AUC < 0.7). As the sequence length increased from 41 to 201, AUC values increased or plateau for specific encodings, especially physicochemical and composition-based encodings (DNC, TNC, CKSNAP, RCKmer, PseEIIP, NCP, and EIIP), indicating that adding sequence context improves prediction accuracy and eventually reaches saturation. Shorter windows (<81nt) often yielded lower AUCs, suggesting insufficient contextual information to capture Ψ-associated sequence patterns. These results demonstrate that increasing sequence length effectively capture Ψ-related information and enhances performance up to a moderate range. In addition, correlation-based encodings such as PseKNC, PseDNC, and SCPseDNC exhibited an upward trend with increasing sequence length, while their AUCs were not high.

**Figure 3.**
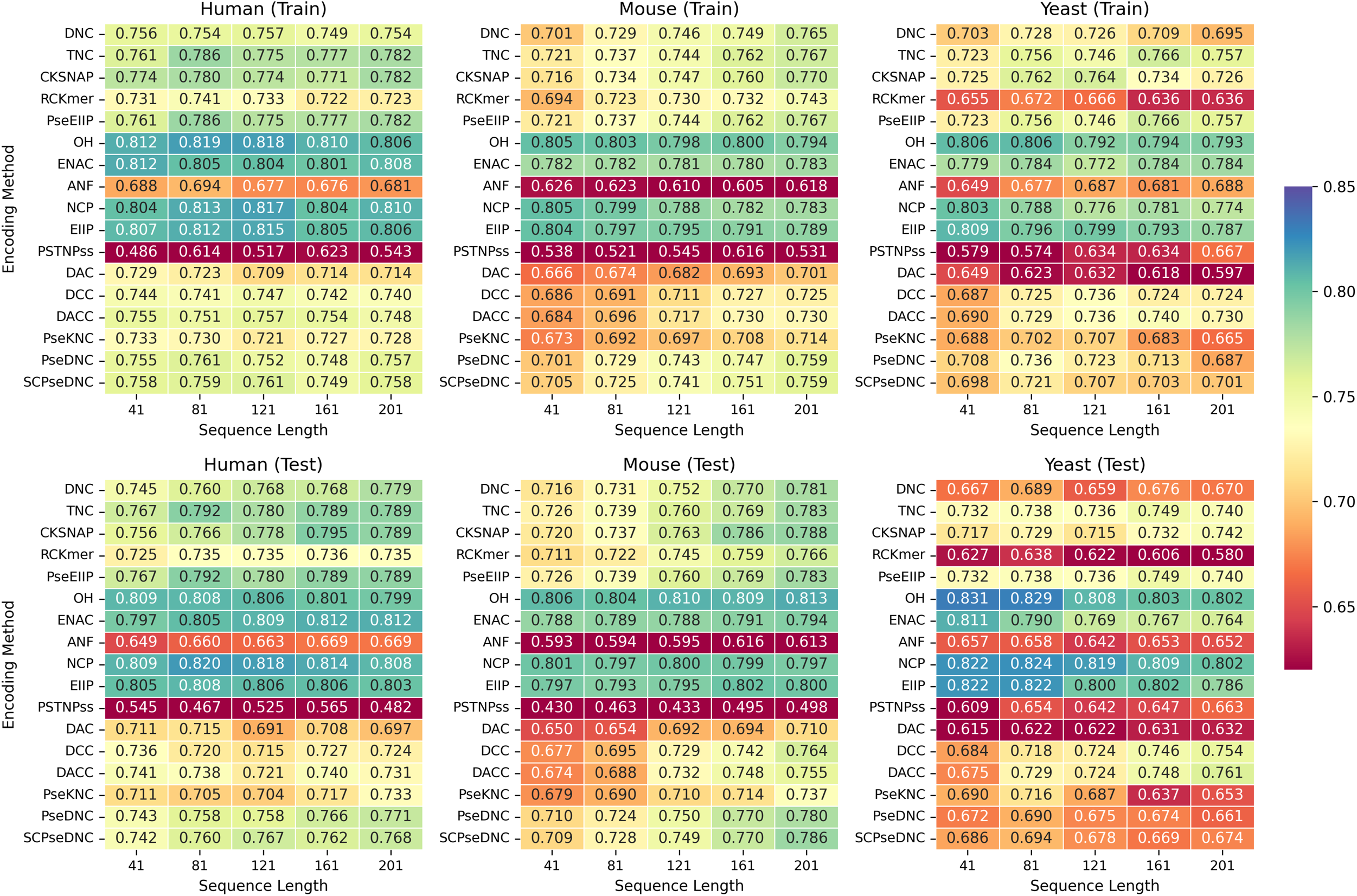
Comparative AUC performance of LGBM-based baseline models using 17 encoding methods with respect to varying sequence lengths in human, mouse, and yeast. The upper and lower panels show the AUCs on the training and test dataset, respectively

### 3.3 DL performance trends with varying sequence lengths

The AUCs for CNN, BiLSTM, and TX models with two encoding schemes (W2V and OH) were computed on the training and test datasets at five sequence lengths (41, 81, 121, 161, and 201) in three species of human, mouse, and yeast, as shown in Figure 4. Across all datasets, the CNN models achieved the highest AUCs on both training and test datasets. In contrast, BiLSTM models presented a downward trend in AUC with increasing sequence length. For the human dataset, CNN-W2V achieved AUCs of 0.830-0.844 on the training set and 0.828-0.857 on the test dataset, maintaining substantial generalization. CNN yielded very high AUCs at moderate sequence lengths (81–121) on both the training and test datasets. Shorter sequence of 41 provided limited context, resulting in slightly lower predictive performance, whereas very long sequences (≥161) tended to degrade AUCs. The BiLSTM–W2V model, however, dropped from AUC of 0.771 in training to 0.756 in testing at a length of 41 in human, and further declined below 0.657 at longer lengths (≥161), suggesting low generalizability and vanishing gradient effects in modeling long dependencies. Similar trends were observed in the mouse and yeast datasets, indicating difficulty in effectively exploiting long-range representations. TX models exhibited intermediate performances between CNN and BiLSTM.

**Figure 4.**
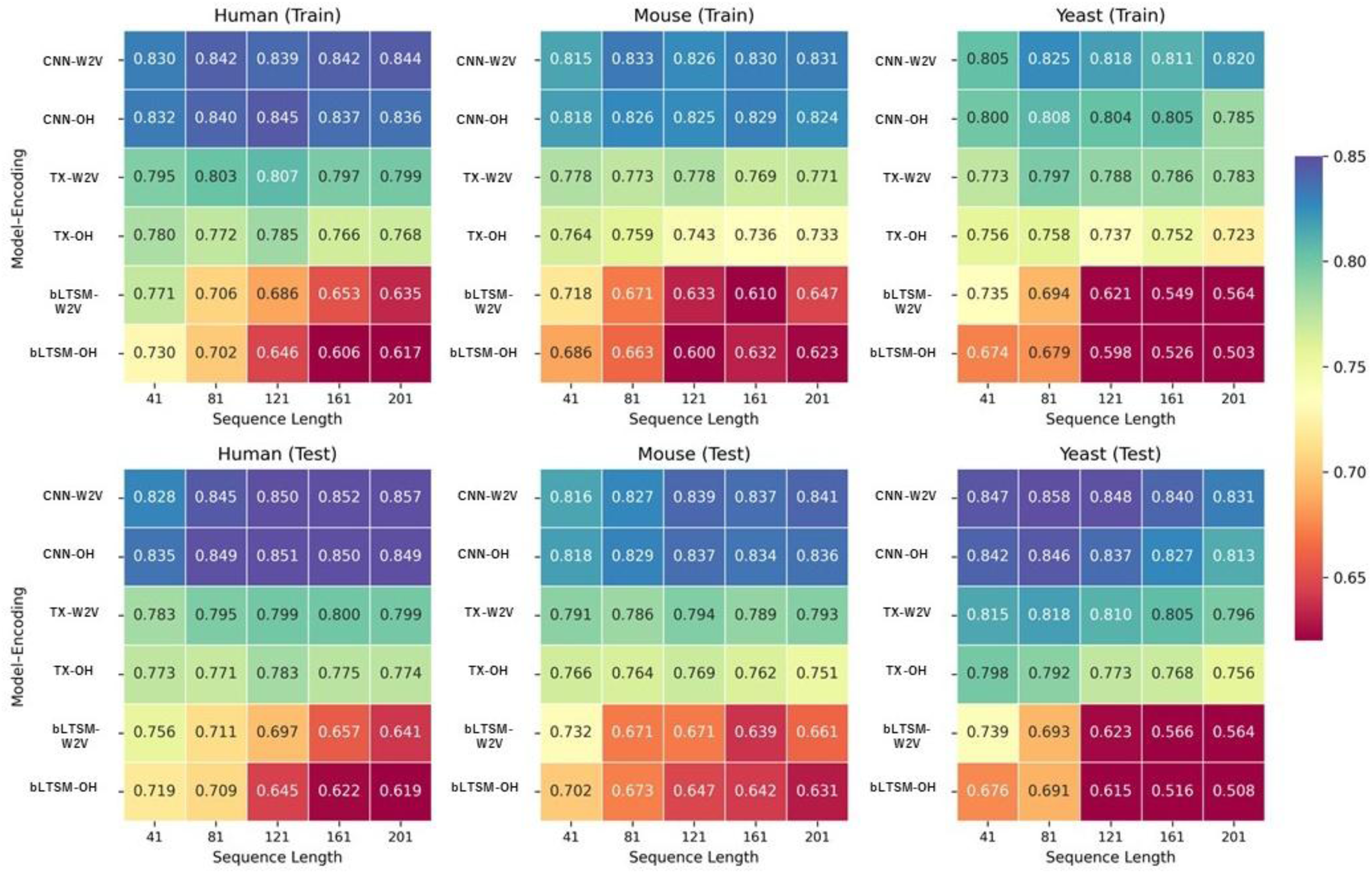
Comparative AUC performance of CNN-based baseline models using W2V and OH encoding methods with respect to varying sequence lengths in human, mouse, and yeast. The upper and lower panels show the AUCs on the training and test dataset, respectively

Across all datasets, W2V encoding consistently outperformed OH encoding for every architecture. The performance gap between W2V and OH was most notable in BiLSTM and TX models, underscoring the advantage of dense, context-aware embeddings in capturing sequence semantics. In the human test data, CNN–W2V achieved an AUC of 0.857 at a length of 201, compared with 0.849 for CNN-OH, while TX-W2V reached 0.800 versus 0.775 for TX-OH at a length of 161; BiLSTM-W2V reached 0.756 versus 0.719 for BiLSTM-OH at a length of 41. This trend suggests that distributed representations provided by the W2V embeddings are more effective in sequence-based learning than the sparse OH.

### 3.4. Meta-PseU construction

To construct a meta-classifier model, we concatenated the predicted probability scores produced by multiple baseline models with a sequence window of 201 on the training dataset and input them into an LR model, as shown in Figure 1. The LR-based stacking models were trained using a 5-fold CV. Concretely, we evaluated the AUC values for all 125 baseline models on the training dataset and added the baseline models in the descending order of the AUCs. Across all datasets, the AUC of the stacking models tended to improve as more baseline models were added; the performance gains became saturated beyond a certain point. Based on these results, the configuration at which performance improvement saturated was defined as the optimal number of the baseline models. In this study, we selected 32 baseline models based on their AUC rankings (**Tables S3, S4, S5**). DL models, including CNN-W2V and CNN-OH, were among the highest-ranked methods. The ML models employing physicochemical encodings (EIIP, and NCP) as well as composition-based encodings (TNC, PseEIIP, and CKSNAP) consistently achieved high rankings across all datasets.

### 3.5. Optimal sequence length of Meta-PseU

We assessed how sequence length affects the prediction performance of the stacking model consisting of 32 baseline models on the training datasets of human, mouse, and yeast, as shown in Figure 5. Overall, extending the sequence length consistently improved AUC across datasets, indicating that longer sequences provide more abundant contextual information for the ensemble. The AUC plateaued beyond a certain length, suggesting diminishing returns with excessive context. Consequently, the optimal sequence lengths (those yielding the highest AUC) for human, mouse and yeast were 201, 161, and 161, respectively.

**Figure 5.**
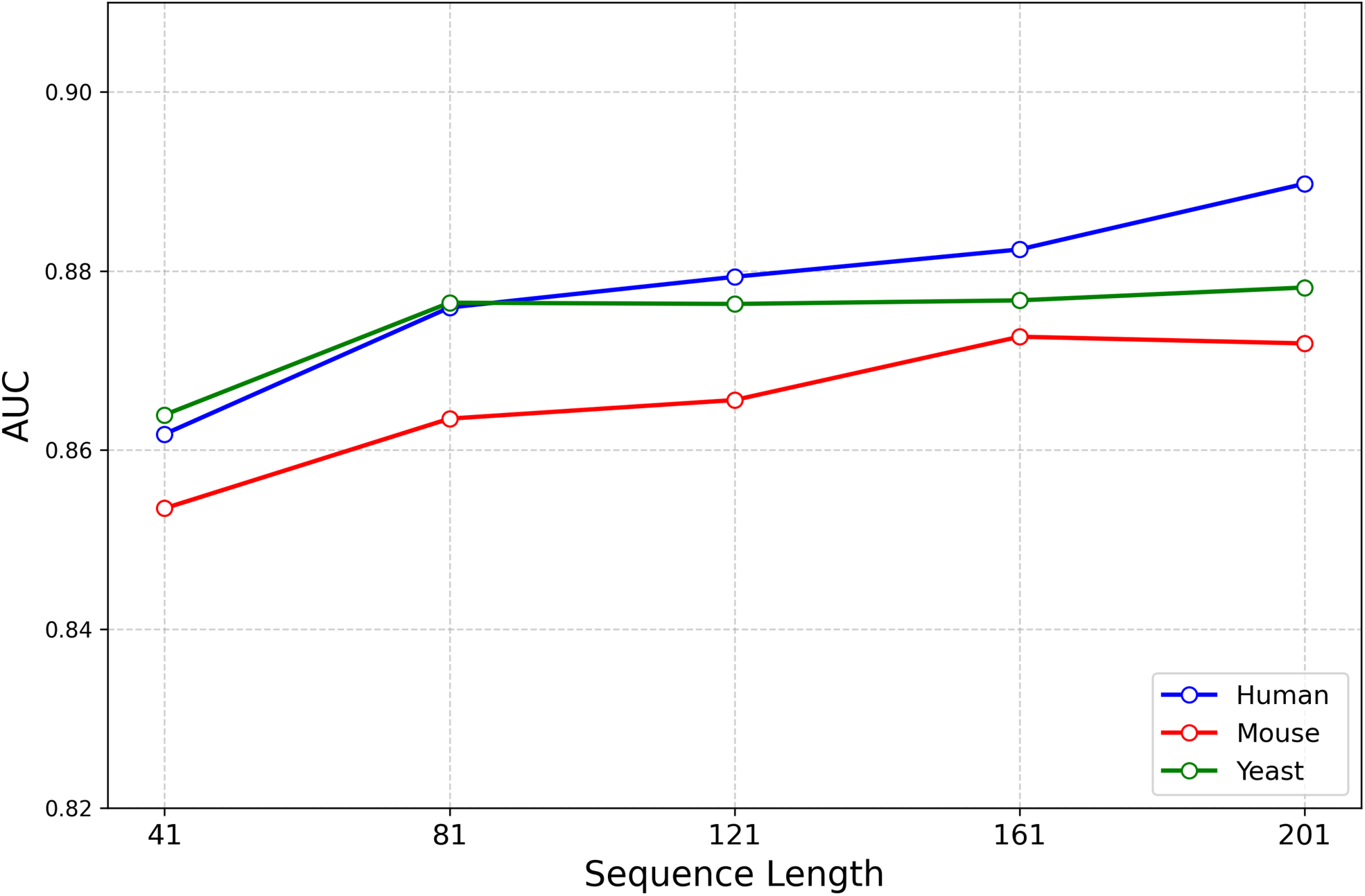
Changes in AUC with increasing sequence length for Meta-PseU on the training datasets of human, mouse and yeast.

### 3.6. Comparison of Meta-PseU with SOTAs

To evaluate the effectiveness of Meta-PseU, we compared Meta-PseU with two SOTA methods (PseU-ST and RSCNN-PseU) with an optimal sequence length on the training and test datasets, as shown in **Tables 2** and **3**. PseU-FKeERF was excluded from the comparison due to its extremely high computational cost, which made comprehensive evaluation across all datasets impractical. In the training datasets, RSCNN-PseU achieved the highest performance across all species (human, mouse, and yeast), with AUC values of 0.940–0.950 and MCC values of 0.785–0.845. In contrast, Meta-PseU demonstrated the second-highest performance while clearly outperforming the conventional method PseU-ST. Notably, for AUC, PseU-ST achieved 0.807 for human, 0.815 for mouse, and 0.844 for yeast, whereas Meta-PseU achieved higher AUC values than PseU-ST in all species datasets.

**Table 2.** Comparison of Meta-PseU stacking models with the two SOTA models with the optimal sequence length on the training datasets.

| Species | Length | Model | SEN | SPE | ACC | MCC | AUC |
| --- | --- | --- | --- | --- | --- | --- | --- |
| Human | 201 | Meta-PseU | 0.735 | **0.889** | 0.812 | 0.631 | 0.890 |
| Human | 201 | PseU-ST | 0.693 | 0.759 | 0.726 | 0.452 | 0.807 |
| Human | 201 | RSCNN-PseU | **0.907** | 0.877 | **0.892** | **0.785** | **0.94** |
| Mouse | 161 | Meta-PseU | 0.835 | 0.764 | 0.8 | 0.603 | 0.878 |
| Mouse | 161 | PseU-ST | 0.740 | 0.724 | 0.732 | 0.464 | 0.815 |
| Mouse | 161 | RSCNN-PseU | **0.911** | **0.912** | **0.911** | **0.823** | **0.947** |
| Yeast | 161 | Meta-PseU | 0.764 | 0.871 | 0.818 | 0.643 | 0.892 |
| Yeast | 161 | PseU-ST | 0.729 | 0.801 | 0.765 | 0.531 | 0.844 |
| Yeast | 161 | RSCNN-PseU | **0.933** | **0.912** | **0.922** | **0.845** | **0.95** |

**Table 3.** Comparison of Meta-PseU stacking models with the two SOTA models with the optimal sequence length on the test datasets.

| Species | Length | Model | SEN | SPE | ACC | MCC | AUC |
| --- | --- | --- | --- | --- | --- | --- | --- |
| Human | 201 | Meta-PseU | **0.719** | **0.873** | **0.796** | **0.599** | **0.874** |
| Human | 201 | PseU-ST | 0.683 | 0.732 | 0.707 | 0.415 | 0.760 |
| Human | 201 | RSCNN-PseU | 0.629 | 0.566 | 0.597 | 0.195 | 0.640 |
| Mouse | 161 | Meta-PseU | **0.825** | **0.746** | **0.786** | **0.577** | **0.871** |
| Mouse | 161 | PseU-ST | 0.702 | 0.691 | 0.697 | 0.393 | 0.765 |
| Mouse | 161 | RSCNN-PseU | 0.555 | 0.58 | 0.567 | 0.135 | 0.598 |
| Yeast | 161 | Meta-PseU | **0.728** | **0.853** | **0.791** | **0.589** | **0.877** |
| Yeast | 161 | PseU-ST | 0.660 | 0.723 | 0.692 | 0.384 | 0.783 |
| Yeast | 161 | RSCNN-PseU | 0.663 | 0.575 | 0.619 | 0.239 | 0.664 |

However, the performance trend changed dramatically in the independent test datasets. RSCNN-PseU exhibited substantial drops in SEN, SPE, and AUC. The AUC fell to 0.640, 0.598, and 0.664 for human, mouse and yeast, respectively, indicating serious overfitting. In contrast, Meta-PseU exhibited relatively consistent performance between the training and independent test datasets. Specifically, Meta-PseU achieved AUCs of 0.890 and 0.874 on the human training and test sets, respectively; 0.878 and 0.871 in mouse; and 0.892 and 0.877 in yeast. These results indicates that Meta-PseU possesses more generalization capability than PseU-ST and RSCNN-PseU across the three species and maintains robust predictive accuracy even for previously unseen RNA sequences. Across all species, Meta-PseU consistently outperformed the SOTA models. It achieved 15% and 37% higher AUC than PseU-ST and RSCNN-PseU on the human dataset;14% and 46% higher on the mouse dataset; and 12% and 32% higher on the yeast dataset.

### 3.7. Shapley additive explanation (SHAP) analysis

To interpret some mechanisms underlying Meta-PseU, we performed SHAP analysis on the probability scores generated by the baseline models incorporated into the framework [52, 53] (Figure 6). SHAP values were computed for the human, mouse, and yeast datasets to quantify the contribution of each baseline model to the final prediction output. As a result, probabilistic features derived from CNN- and LGBM-based models consistently exhibited strong influence across all species, suggesting that contextual sequence representations captured by CNNs and feature optimization achieved by boosting-based models play central roles in driving Meta-PseU’s predictions. In human, CNN-W2V and CNN-OH contributed the most, followed by LGBM-OH, LGBM-NCP, LGBM-EIIP, and LGBM-TNC. In mouse and yeast, CNN-W2V and CNN-OH also ranked highly. As expected, the physicochemical property and composition-based encodings such as NCP, EIIP, and TNC, which presented high prediction performance or an upward trend with increasing sequence length, were observed among the top-ranked methods across all the species. These findings indicate that Meta-PseU effectively learns complementary features from diverse encodings and classifiers.

**Figure 6.**
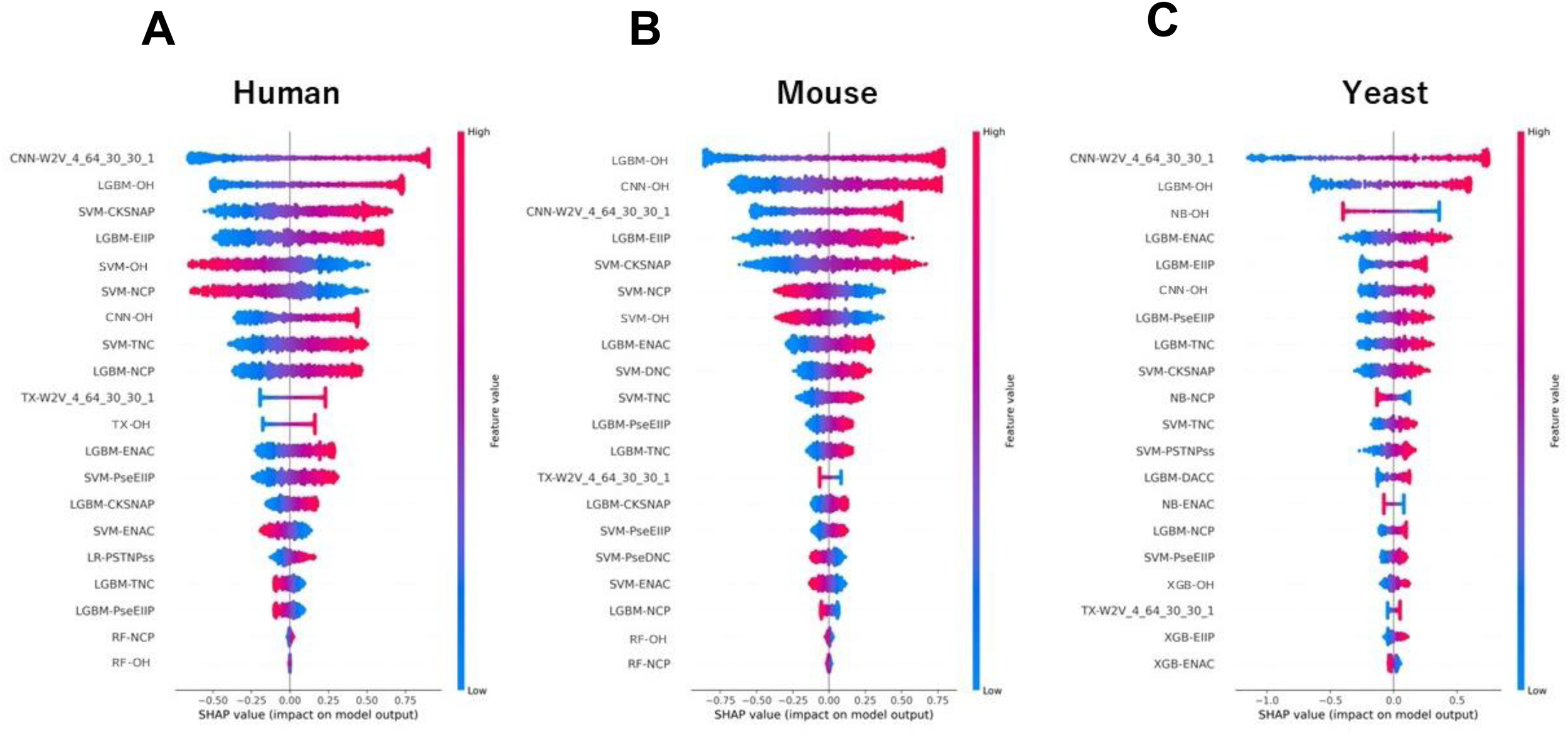
SHAP summary plots showing model-level contributions for human (A), mouse (B), and yeast (C). This figure visualizes the contribution of baseline models to the prediction output using SHAP values. Each point represents a sample, where the horizontal axis indicates the impact on model output (positive or negative contribution). Colors reflect the feature value (red = high, blue = low). Models are ranked vertically by their overall importance. These plots help identify which models are consistently influential across species and which are species-specific.

### 3.8. UMAP-based visualization of probabilistic feature representations

To visualize whether the probabilistic feature representations learned by Meta-PseU preserve discriminative capability across species, we applied UMAP to the 32-dimensional probability vectors obtained from the baseline models included in the stacking framework [54] (Figure 7). In both the human and mouse test datasets, positive and negative samples formed clearly separated clusters. These findings indicate that the probabilistic feature representations maintain a stable and reproducible decision boundary even for unseen data, demonstrating strong generalization capability. In contrast, the yeast dataset exhibited more dispersed clustering compared with human and mouse, reflecting species-specific sequence characteristics. Meta-PseU was capable of effectively capturing biologically meaningful modification signatures across three species.

**Figure 7.**
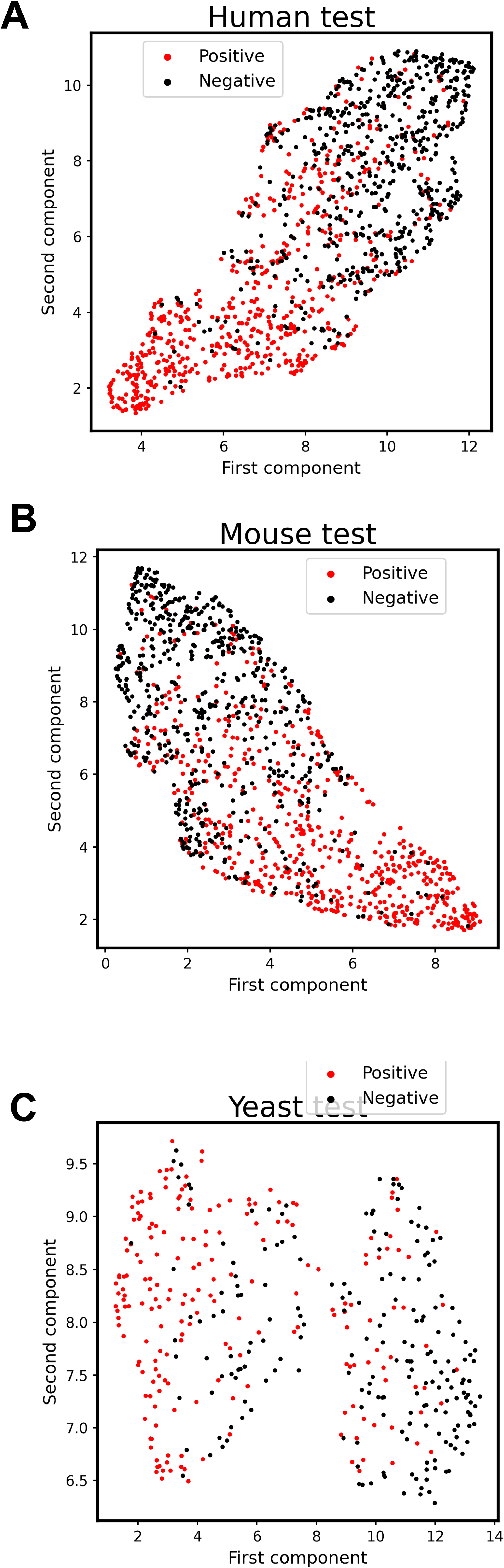
UMAP projection of stacking probabilistic score vectors on the test datasets of human (A), mouse (B), and yeast (C). Red points indicate positive (modified) samples, and black points indicate negative samples. UMAP preserves both global and local structure; therefore, the degree of cluster separation reflects how well the encoding captures the intrinsic differences between positive and negative samples.

## 4. Discussion and conclusions

In Ψ site identification, computational prediction has become indispensable for complementing experimental identification. However, existing ML and DL predictors suffer from major limitations, including small and inconsistent datasets, poor generalizability across species, and limited reproducibility (**Tables S1**, **S2**, **Table 3**). To overcome these issues, we constructed new long-sequence datasets derived from RMBase 3.0 and developed a novel computational Ψ-site predictor, Meta-PseU. It considered 125 baseline models including seven ML algorithms with 17 encoding methods and three DL models based on W2V embeddings. The probabilistic outputs of these models were combined using LR as a meta-classifier in a stacking framework to produce the final predictions. By optimizing model configuration, Meta-PseU stacking 32 baseline models achieved the best performance on the test datasets of all the species. Meta-PseU achieves higher accuracy and significantly improved prediction performance compared with the SOTAs across human, mouse, and yeast.

In contrast to previous studies that relied on short 21-necleotide windows, we analyzed the performance of all the baseline models with varying sequence lengths from 41 to 201. As the sequence length increases, composition-based and physicochemical property-based encodings such as DNC, TNC, CKSNAP, RCKmer, PseEIIP, NCP, and EIIP tended to increase AUCs. This trend is caused by the extended windows capturing broader contextual dependencies. For instance, regional GC content, repetitive motifs, or structural signals may influence pseudouridylation. Such contextual enrichment cannot be adequately represented in very short windows. In addition, correlation-based encodings such as PseKNC, PseDNC, and SCPseDNC exhibited an upward trend with increasing sequence length, although their AUCs were not high. They rely on representing both local and global sequence-order information through statistical correlations between nucleotide pairs. These methods capture how nucleotides separated by different distances interact, providing insight into long-range sequence dependencies that may influence Ψ formation. When the sequence window is short, these correlations are not easy to estimate, resulting in unstable and less discriminative features. As the sequence length increases, more nucleotide pairs become available, enabling more reliable estimation of these correlation factors and improving feature robustness. Longer sequences also provide additional contextual information, allowing the model to capture distant dependencies related to RNA secondary structure or enzyme recognition. Furthermore, the CNN models indicated higher AUCs across species and positive effects of increasing sequence lengths, confirming their capability to extract local motifs and their distant relationship. On the other hand, both BiLSTM and TX models reduced AUCs for long sequences, likely due to gradient instability and attention dilution effects, respectively.

Finally, we benchmarked Meta-PseU against the SOTA predictors, including PseU-ST and RSCNN-PseU. The SOTAs achieved high performance on training datasets for human, mouse and yeast, but they failed to generalize to independent test datasets. In contrast, Meta-PseU substantially reduced the performance gaps between training and independent test datasets. Meta-PseU suppressed such overfitting and demonstrated superior generalization across all the species. Furthermore, our newly constructed long-sequence datasets establish a new benchmark for RNA Ψ site prediction and provide a robust foundation for future studies of Ψ biology and disease-associated RNA regulation.

## Data and Code Availability

The python programs of Meta-PseU including training and testing processes and all the datasets are freely accessible at https://github.com/kuratahiroyuki/MetaPseU. Any additional information required to reproduce or further analyze the results presented in this study is available upon reasonable request to the corresponding author.

## Author Contributions

**Takumi Sutou**: Methodology, programming, analysis, writing. **Md. HARUN-OR-ROSHID**, Methodology and programming. **Hiroyuki Kurata**: Supervision, conceptualization, project administration, data curation, funding acquisition, and writing.

## Declarations

While preparing this work, the authors used ChatGPT based on the GPT-5.1 architecture to improve the readability. After using this tool, the authors reviewed and edited the contents as needed and take full responsibility for the content s.

## Ethics statement

The authors declare that the current study did not involve any experiments on humans or animals. All computational analyses were conducted using publicly available data and software tools, and no ethical approval was required. The research complies with the ethical standards of scientific integrity, transparency, and reproducibility. All authors have read and approved the manuscript and agree with its submission.

