## Supplemental Information for "Meta-PseU: A Meta-Classifier for Robust Prediction of RNA Pseudouridine Modification Sites from Long Sequences"

Table S1. Summary of existing Ψ-site prediction models with a sequence length of 21 in human, mouse and yeast. “Reported performance” means the reported values in the original papers; “Reproduced performance” indicates our simulated values in this study.

| Model | Server | Program | Species | Reported Performance | Comments with our reproduced performance |
|---|---|---|---|---|---|
| RSCNN-PseU (2025) | × | ○ | *H. sapiens*<br>*S. cerevisiae*<br>*M. musculus* | Training<br>AUC in human: 0.9511<br>AUC in mouse: 0.9552<br>AUC in yeast: 0.9622<br>Independent test<br>AUC in human: 0.978<br>AUC in mouse: N/A<br>AUC in yeast: 0.968 | Overfitting was observed in the human and yeast datasets.<br>Reproduced independent test<br>AUC in human: 0.6707<br>AUC in mouse: NA<br>AUC in yeast: 0.5725 |
| PseU-FKeERF (2024) | × | ○ | *H. sapiens*<br>*S. cerevisiae*<br>*M. musculus* | Training<br>AUC in human: 0.7814<br>AUC in mouse: 0.7784<br>AUC in yeast: 0.8803<br>Independent test<br>AUC in human: 0.9254<br>AUC in mouse: N/A<br>AUC in yeast: 0.9468 | Overfitting occurred in the human dataset.<br>Reproduced independent test<br>AUC in human: 04604<br>AUC in mouse: NA<br>AUC in yeast: 0.8659 |
| PseU-ST (2023) | × | ○ | *H. sapiens*<br>*S. cerevisiae*<br>*M. musculus* | Training<br>AUC in human: 0.9856<br>AUC in mouse: 0.9620<br>AUC in yeast: 0.9595<br>Independent test<br>No result of AUC was documented. | Overfitting was observed in the human and yeast datasets.<br>Reproduced independent test<br>AUC in human: 0.7055<br>AUC in mouse: NA<br>AUC in yeast: 0.7655 |
| iPseU-TWSSVM (2022) | ○ | × | *H. sapiens*<br>*S. cerevisiae*<br>*M. musculus* | Training<br>AUC in human: 0.682<br>AUC in mouse: 0.775<br>AUC in yeast: 0.758<br>Independent test<br>AUC in human: 0.786<br>AUC in mouse: N/A<br>AUC in yeast: 0.905 | Web server<br>http://bioinformatics.hitsz.edu.cn/Pse-in-One2.0/<br>is not available |
| PorPoise | ○ | × | *H. sapiens* | Training | Web server |

| | | | | | |
|---|---|---|---|---|---|
| (2021) | | | *S. cerevisiae*<br>*M. musculus* | AUC in human: 0.733<br>AUC in mouse: 0.737<br>AUC in yeast: 0.768<br>Independent test<br>No result of AUC was documented. | http://web.unimelb-bioinfortools.cloud.edu.au/Porpoise/ is not available |
| PseUdeep (2021) | × | ○ | *H. sapiens*<br>*S. cerevisiae*<br>*M. musculus* | Training<br>AUC in human: 0.74<br>AUC in mouse: 0.77<br>AUC in yeast: 0.74<br>Independent test<br>AUC in human: 0.720<br>AUC in mouse: N/A<br>AUC in yeast: 0.909 | Program does not implement any independent test process. Overfitting was observed in the yeast datasets.<br>Reproduced independent test<br>AUC in human: 0.74<br>AUC in yeast: 0.30 |
| EnsemPseU (2020) | × | × | *H. sapiens*<br>*S. cerevisiae*<br>*M. musculus* | Training<br>AUC in human:0.786<br>AUC in mouse: 0.775<br>AUC in yeast: 0.700<br>Independent test<br>No result of AUC was documented. | |
| XG-PseU (2020) | ○ | × | *H. sapiens*<br>*S. cerevisiae*<br>*M. musculus* | Training<br>AUC in human: 0.70<br>AUC in mouse: 0.74<br>AUC in yeast: 0.77<br>Independent test<br>No result of AUC was documented. | Web server<br>http://www.bioml.cn/<br>is not available. |
| iPseU-layer (2020) | × | × | *H. sapiens*<br>*S. cerevisiae*<br>*M. musculus* | The results of the AUC were not documented. | |
| RF-PseU (2020) | ○ | × | *H. sapiens*<br>*S. cerevisiae*<br>*M. musculus* | Training<br>AUC in human: 0.70<br>AUC in mouse: 0.796<br>AUC in yeast: 0.810<br>Independent test<br>AUC in human: 0.800 | Web server<br>http://148.70.81.170:10228/rfpseu.<br>is not available. |

| | | | | | |
|---|---|---|---|---|---|
| | | | | AUC in mouse: N/A<br>AUC in yeast: 0.838 | |
| iPseU-NCP<br>(2019) | ○ | × | *H. sapiens*<br>*S. cerevisiae*<br>*M. musculus* | No result of AUC was documented. | Web server<br>http://45.117.83.253/problem-iPseU-NCP<br>is not available. |
| PseUI<br>(2018) | ○ | × | *H. sapiens*<br>*S. cerevisiae*<br>*M. musculus* | Training<br>AUC in human: 0.68<br>AUC in mouse: 0.77<br>AUC in yeast: 0.69<br>Independent test<br>No result of AUC was documented. | Web server<br>http://zhulab.ahu.edu.cn/PseUI.<br>is not available. |

Table S2 Comparison between the reported and reproduced performances of three SOTAs on the training and test datasets. The length of sequences is 21.

| **Model** | | **5-fold cross-validation** | | | | | **Independent test** | | | | |
|---|---|---|---|---|---|---|---|---|---|---|---|
| | | ACC (%) | MCC (%) | SEN (%) | SPE (%) | AUC (%) | ACC (%) | MCC (%) | SEN (%) | SPE (%) | AUC (%) |
| **PseU-ST(RF+LR)** | Reported | 93.64 | 87.28 | 94.34 | 92.93 | 98.56 | 89.00 | 79.02 | 97.00 | 81.00 | 96.51 |
| | **Reproduced** | 93.56 | 87.28 | 94.34 | 92.93 | 98.56 | **63.50** | **27.16** | **58.00** | **69.00** | **66.69** |
| **RSCNN-PseU** | Reported | 92.53 | 85.17 | 94.31 | 90.78 | 95.11 | 96.50 | 93.02 | 96.00 | 97.00 | 97.80 |
| | **Reproduced** | 93.94 | 88.05 | 93.47 | 94.39 | 95.55 | **63.20** | **26.53** | **58.90** | **67.50** | **67.07** |
| **PseU-FKeERF** | Reported | 78.08 | 56.44 | 77.62 | 78.66 | 78.14 | 99.19 | 85.04 | 90.11 | 94.97 | 92.54 |
| | **Reproduced** | 88.88 | 78.14 | 91.57 | 86.37 | 95.57 | **46.69** | **-0.67** | **43.59** | **49.8** | **46.04** |

"Reported" means the reported values in the original papers; "Reproduced" indicates our simulated values in this study. We reproduced and evaluated the prediction performance of three publicly available state-of-the-art (SOTA) models: PseU-ST, RSCNN-PseU, and PseU-FKeERF. PseU-ST exhibited high prediction performances on the training datasets of human but provided poor reproducibility and generalizability on the independent test dataset. PseU-FKeERF and RSCNN-PseU also exhibited less reproducibility and less generalizability on the independent test datasets. Thus, the reproducibility and generalizability of these models remain to be fully clarified.

Table S3 AUC-based performance ranking of baseline models on the human training dataset

| ML/DL | Encodings | SEN | SPE | ACC | MCC | AUC |
|---|---|---|---|---|---|---|
| **CNN** | W2V | 0.708 | 0.827 | 0.768 | 0.540 | 0.844 |
| **CNN** | OH | 0.664 | 0.849 | 0.756 | 0.523 | 0.836 |
| **LGBM** | OH | 0.662 | 0.820 | 0.741 | 0.494 | 0.814 |
| **LGBM** | NCP | 0.627 | 0.850 | 0.739 | 0.490 | 0.811 |
| **LGBM** | EIIP | 0.565 | 0.892 | 0.729 | 0.486 | 0.808 |
| **LGBM** | ENAC | 0.693 | 0.785 | 0.739 | 0.481 | 0.805 |
| **TX** | W2V | 0.709 | 0.742 | 0.725 | 0.451 | 0.799 |
| **SVM** | CKSNAP | 0.595 | 0.845 | 0.720 | 0.458 | 0.794 |
| **SVM** | TNC | 0.653 | 0.798 | 0.725 | 0.459 | 0.793 |
| **SVM** | PseEIIP | 0.690 | 0.759 | 0.724 | 0.454 | 0.792 |
| **LGBM** | PseEIIP | 0.672 | 0.755 | 0.714 | 0.430 | 0.780 |
| **LGBM** | TNC | 0.672 | 0.755 | 0.714 | 0.430 | 0.780 |
| **LGBM** | CKSNAP | 0.609 | 0.808 | 0.709 | 0.431 | 0.779 |
| **SVM** | ENAC | 0.684 | 0.719 | 0.702 | 0.415 | 0.774 |
| **RF** | OH | 0.594 | 0.795 | 0.695 | 0.405 | 0.772 |
| **LR** | PSTNPss | 0.712 | 0.700 | 0.706 | 0.421 | 0.772 |
| **TX** | OH | 0.739 | 0.658 | 0.699 | 0.400 | 0.768 |
| **RF** | NCP | 0.594 | 0.789 | 0.691 | 0.407 | 0.766 |
| **SVM** | NCP | 0.572 | 0.820 | 0.696 | 0.407 | 0.766 |
| **SVM** | OH | 0.597 | 0.799 | 0.698 | 0.407 | 0.766 |
| **SVM** | DNC | 0.592 | 0.812 | 0.702 | 0.417 | 0.765 |
| **LGBM** | DNC | 0.491 | 0.872 | 0.681 | 0.397 | 0.763 |
| **SVM** | PseDNC | 0.526 | 0.858 | 0.692 | 0.412 | 0.763 |
| **LGBM** | PseDNC | 0.488 | 0.877 | 0.682 | 0.399 | 0.761 |
| **LGBM** | SCPseDNC | 0.589 | 0.785 | 0.687 | 0.398 | 0.760 |
| **SVM** | PSTNPss | 0.570 | 0.832 | 0.701 | 0.421 | 0.759 |
| **RF** | EIIP | 0.635 | 0.724 | 0.680 | 0.375 | 0.753 |
| **LGBM** | DACC | 0.553 | 0.819 | 0.686 | 0.387 | 0.752 |
| **XGB** | NCP | 0.602 | 0.779 | 0.691 | 0.388 | 0.750 |
| **LR** | ENAC | 0.652 | 0.727 | 0.690 | 0.386 | 0.749 |
| **RF** | ENAC | 0.666 | 0.687 | 0.676 | 0.372 | 0.748 |
| **RF** | PseDNC | 0.451 | 0.894 | 0.672 | 0.389 | 0.746 |

Table S4 AUC-based performance ranking of baseline models on the mouse training dataset

| ML/DL | Encodings | SEN | SPE | ACC | MCC | AUC |
|---|---|---|---|---|---|---|
| **CNN** | W2V | 0.696 | 0.792 | 0.744 | 0.497 | 0.831 |
| **CNN** | OH | 0.694 | 0.787 | 0.740 | 0.486 | 0.824 |
| **LGBM** | OH | 0.733 | 0.710 | 0.721 | 0.449 | 0.790 |
| **LGBM** | NCP | 0.691 | 0.756 | 0.724 | 0.451 | 0.789 |
| **SVM** | TNC | 0.681 | 0.744 | 0.712 | 0.429 | 0.782 |
| **SVM** | PseEIIP | 0.721 | 0.703 | 0.712 | 0.429 | 0.782 |
| **LGBM** | EIIP | 0.728 | 0.714 | 0.721 | 0.444 | 0.781 |
| **LGBM** | ENAC | 0.685 | 0.741 | 0.713 | 0.430 | 0.778 |
| **SVM** | CKSNAP | 0.598 | 0.817 | 0.708 | 0.426 | 0.773 |
| **LGBM** | CKSNAP | 0.603 | 0.806 | 0.704 | 0.419 | 0.771 |
| **TX** | W2V | 0.689 | 0.694 | 0.691 | 0.383 | 0.771 |
| **SVM** | DNC | 0.637 | 0.770 | 0.704 | 0.415 | 0.768 |
| **LGBM** | PseEIIP | 0.703 | 0.693 | 0.698 | 0.409 | 0.768 |
| **LGBM** | TNC | 0.703 | 0.693 | 0.698 | 0.409 | 0.768 |
| **SVM** | PseDNC | 0.590 | 0.800 | 0.695 | 0.402 | 0.767 |
| **RF** | NCP | 0.681 | 0.701 | 0.691 | 0.390 | 0.764 |
| **RF** | OH | 0.640 | 0.752 | 0.696 | 0.399 | 0.762 |
| **LGBM** | DNC | 0.532 | 0.841 | 0.687 | 0.398 | 0.761 |
| **LGBM** | PseDNC | 0.552 | 0.821 | 0.687 | 0.391 | 0.757 |
| **LGBM** | SCPseDNC | 0.567 | 0.811 | 0.689 | 0.392 | 0.757 |
| **SVM** | NCP | 0.642 | 0.754 | 0.698 | 0.400 | 0.753 |
| **SVM** | OH | 0.640 | 0.755 | 0.697 | 0.399 | 0.753 |
| **RF** | SCPseDNC | 0.477 | 0.880 | 0.679 | 0.391 | 0.750 |
| **LR** | PSTNPss | 0.638 | 0.746 | 0.692 | 0.388 | 0.748 |
| **RF** | DNC | 0.480 | 0.872 | 0.676 | 0.386 | 0.748 |
| **SVM** | ENAC | 0.675 | 0.709 | 0.692 | 0.388 | 0.748 |
| **SVM** | RCKmer | 0.503 | 0.860 | 0.682 | 0.391 | 0.748 |
| **RF** | PseDNC | 0.488 | 0.873 | 0.680 | 0.394 | 0.748 |
| **XGB** | DNC | 0.525 | 0.825 | 0.675 | 0.379 | 0.745 |
| **XGB** | SCPseDNC | 0.554 | 0.806 | 0.680 | 0.380 | 0.745 |
| **XGB** | PseDNC | 0.432 | 0.898 | 0.665 | 0.375 | 0.744 |
| **XGB** | CKSNAP | 0.542 | 0.814 | 0.678 | 0.379 | 0.743 |

Table S6 AUC-based performance ranking of baseline models on the yeast training dataset

| ML/DL | Encodings | SEN | SPE | ACC | MCC | AUC |
|---|---|---|---|---|---|---|
| **CNN** | W2V | 0.777 | 0.678 | 0.728 | 0.458 | 0.820 |
| **LGBM** | OH | 0.574 | 0.887 | 0.731 | 0.491 | 0.793 |
| **CNN** | OH | 0.704 | 0.719 | 0.712 | 0.426 | 0.785 |
| **TX** | W2V | 0.712 | 0.699 | 0.706 | 0.412 | 0.783 |
| **LGBM** | ENAC | 0.616 | 0.830 | 0.723 | 0.461 | 0.781 |
| **LGBM** | NCP | 0.524 | 0.920 | 0.722 | 0.486 | 0.778 |
| **LGBM** | EIIP | 0.526 | 0.904 | 0.715 | 0.465 | 0.777 |
| **SVM** | TNC | 0.796 | 0.653 | 0.724 | 0.455 | 0.773 |
| **SVM** | PseEIIP | 0.797 | 0.639 | 0.718 | 0.445 | 0.770 |
| **LGBM** | TNC | 0.719 | 0.712 | 0.715 | 0.439 | 0.759 |
| **LGBM** | PseEIIP | 0.719 | 0.712 | 0.715 | 0.439 | 0.759 |
| **SVM** | OH | 0.555 | 0.830 | 0.692 | 0.404 | 0.751 |
| **SVM** | NCP | 0.601 | 0.796 | 0.698 | 0.405 | 0.751 |
| **RF** | OH | 0.546 | 0.831 | 0.689 | 0.412 | 0.751 |
| **NB** | NCP | 0.602 | 0.791 | 0.696 | 0.409 | 0.750 |
| **RF** | ENAC | 0.686 | 0.725 | 0.706 | 0.413 | 0.749 |
| **NB** | OH | 0.563 | 0.820 | 0.692 | 0.400 | 0.747 |
| **RF** | EIIP | 0.601 | 0.784 | 0.692 | 0.399 | 0.742 |
| **RF** | TNC | 0.701 | 0.690 | 0.695 | 0.394 | 0.741 |
| **RF** | PseEIIP | 0.701 | 0.690 | 0.695 | 0.394 | 0.740 |
| **NB** | ENAC | 0.690 | 0.691 | 0.690 | 0.383 | 0.740 |
| **XGB** | OH | 0.438 | 0.904 | 0.671 | 0.389 | 0.739 |
| **SVM** | CKSNAP | 0.715 | 0.635 | 0.675 | 0.388 | 0.738 |
| **RF** | NCP | 0.500 | 0.858 | 0.679 | 0.397 | 0.736 |
| **SVM** | PSTNPss | 0.551 | 0.833 | 0.692 | 0.408 | 0.735 |
| **XGB** | ENAC | 0.575 | 0.774 | 0.675 | 0.367 | 0.733 |
| **LR** | PSTNPss | 0.489 | 0.887 | 0.688 | 0.418 | 0.732 |
| **LGBM** | DACC | 0.729 | 0.619 | 0.674 | 0.364 | 0.730 |
| **SVM** | ENAC | 0.666 | 0.692 | 0.679 | 0.370 | 0.730 |
| **XGB** | EIIP | 0.471 | 0.888 | 0.679 | 0.405 | 0.729 |
| **LGBM** | DCC | 0.681 | 0.652 | 0.667 | 0.352 | 0.726 |
| **RF** | PSTNPss | 0.523 | 0.841 | 0.682 | 0.390 | 0.726 |

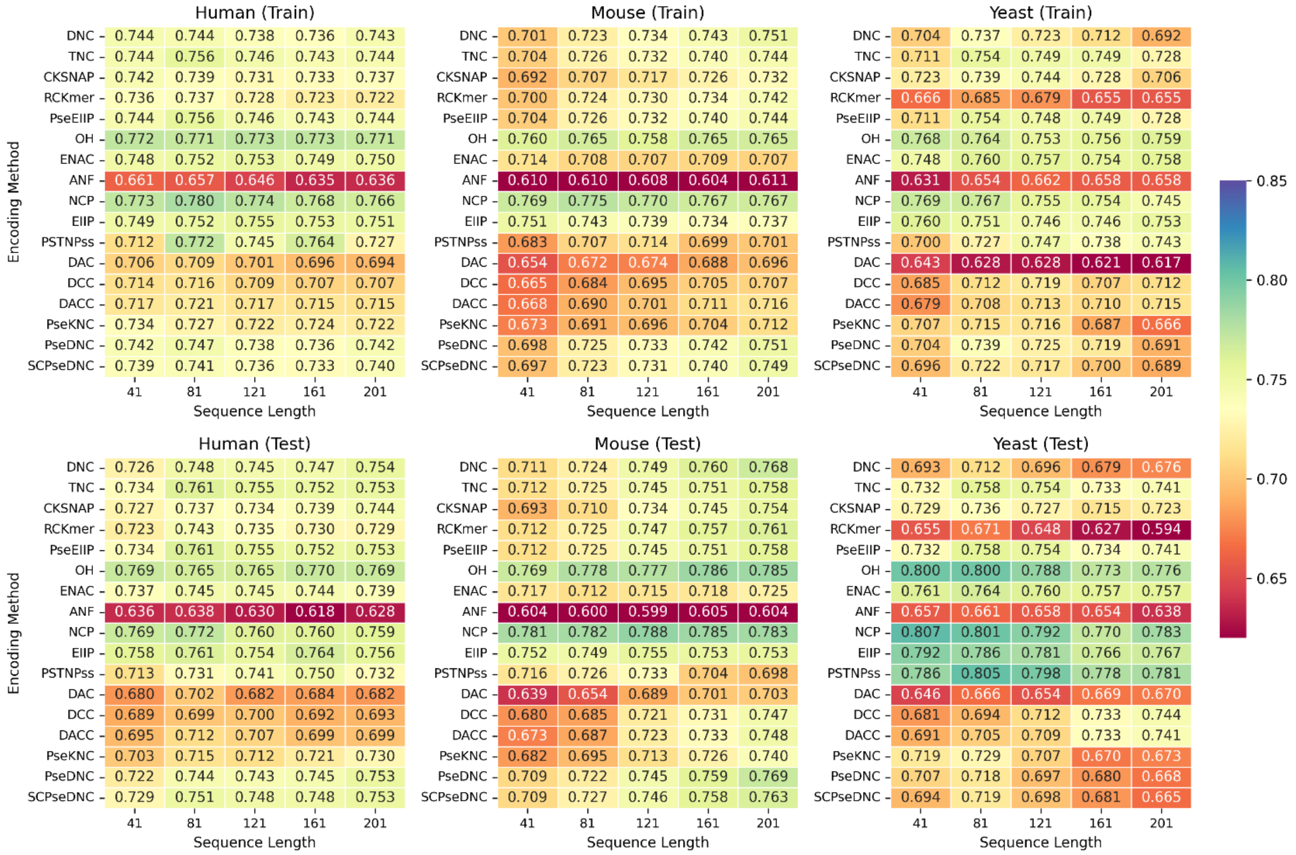


Figure S1 Comparative AUC performance of RF baseline models using 17 encoding methods across varying sequence lengths in human, mouse, and yeast.

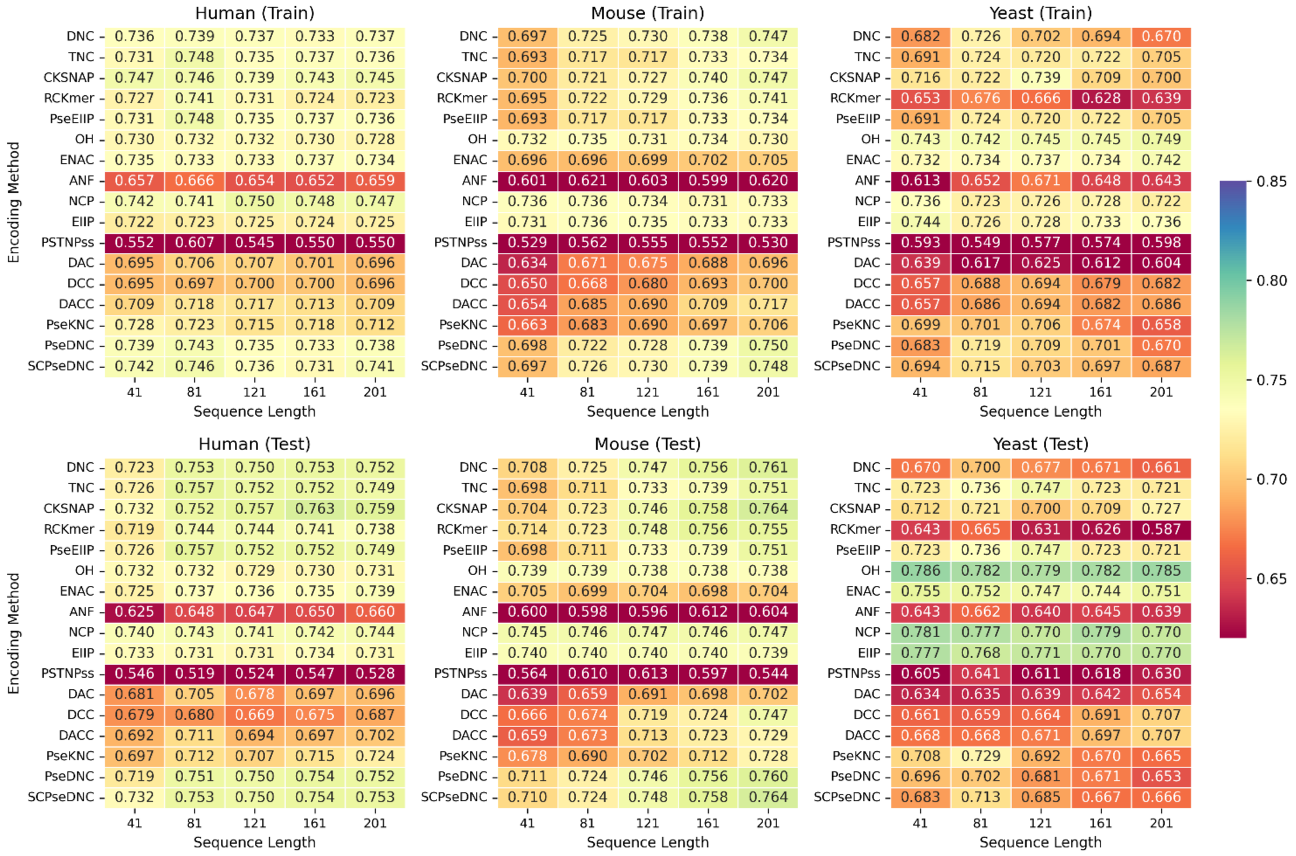

Figure S2 Comparative AUC performance of XGB baseline models using 17 encoding methods across varying sequence lengths in human, mouse, and yeast.

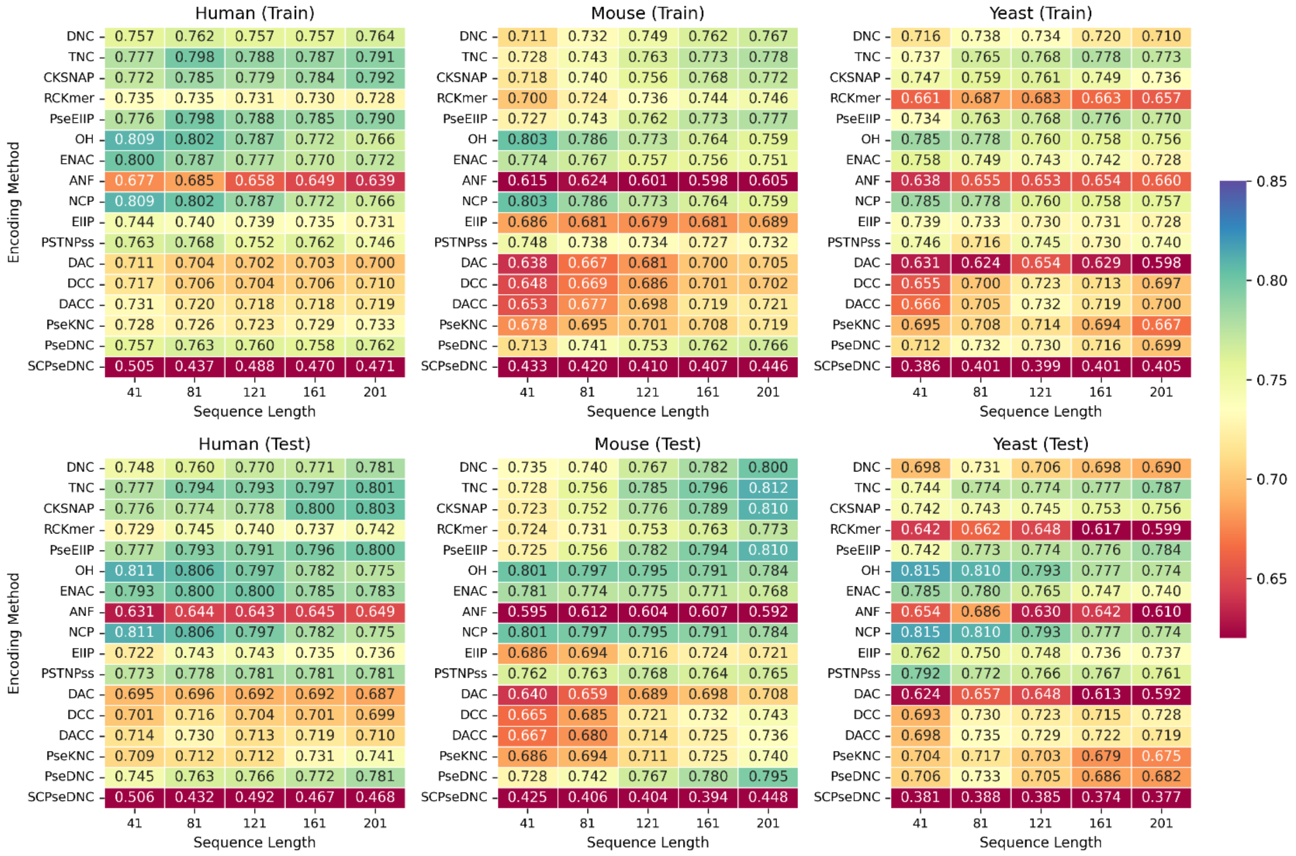


Figure S3 Comparative AUC performance of SVM baseline models using 17 encoding methods across varying sequence lengths in human, mouse, and yeast.

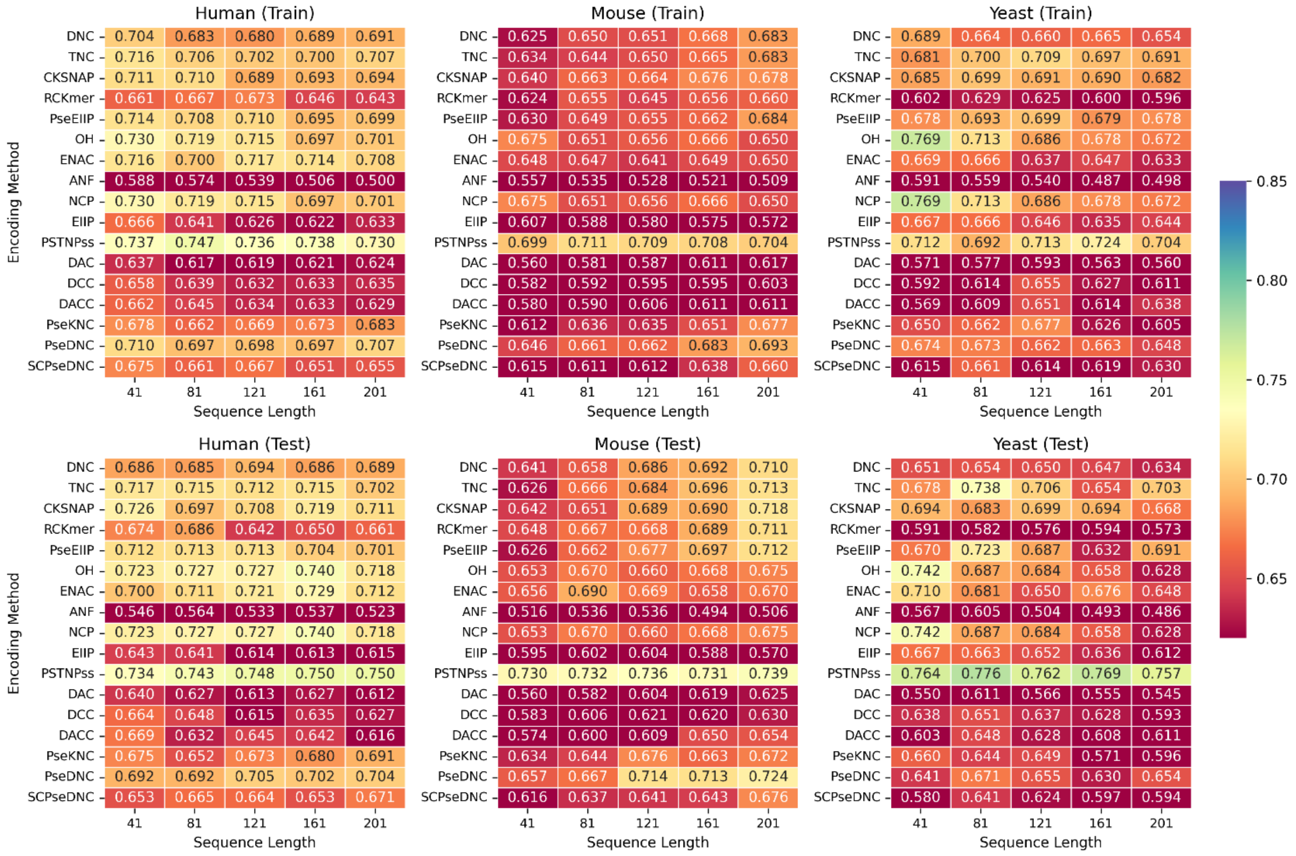


Figure S4 Comparative AUC performance of KNN baseline models using 17 encoding methods across varying sequence lengths in human, mouse, and yeast.

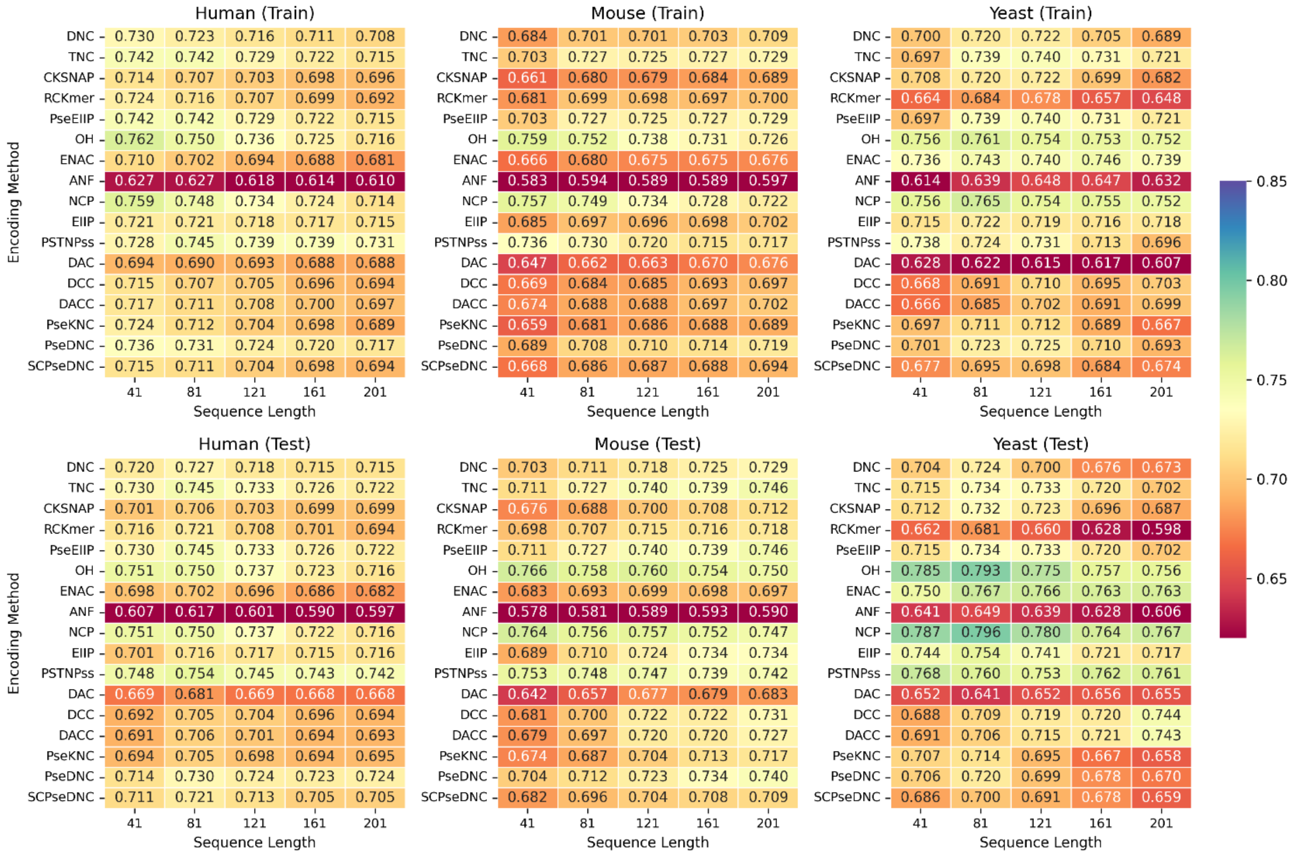


Figure S5 Comparative AUC performance of NB baseline models using 17 encoding methods across varying sequence lengths in human, mouse, and yeast.

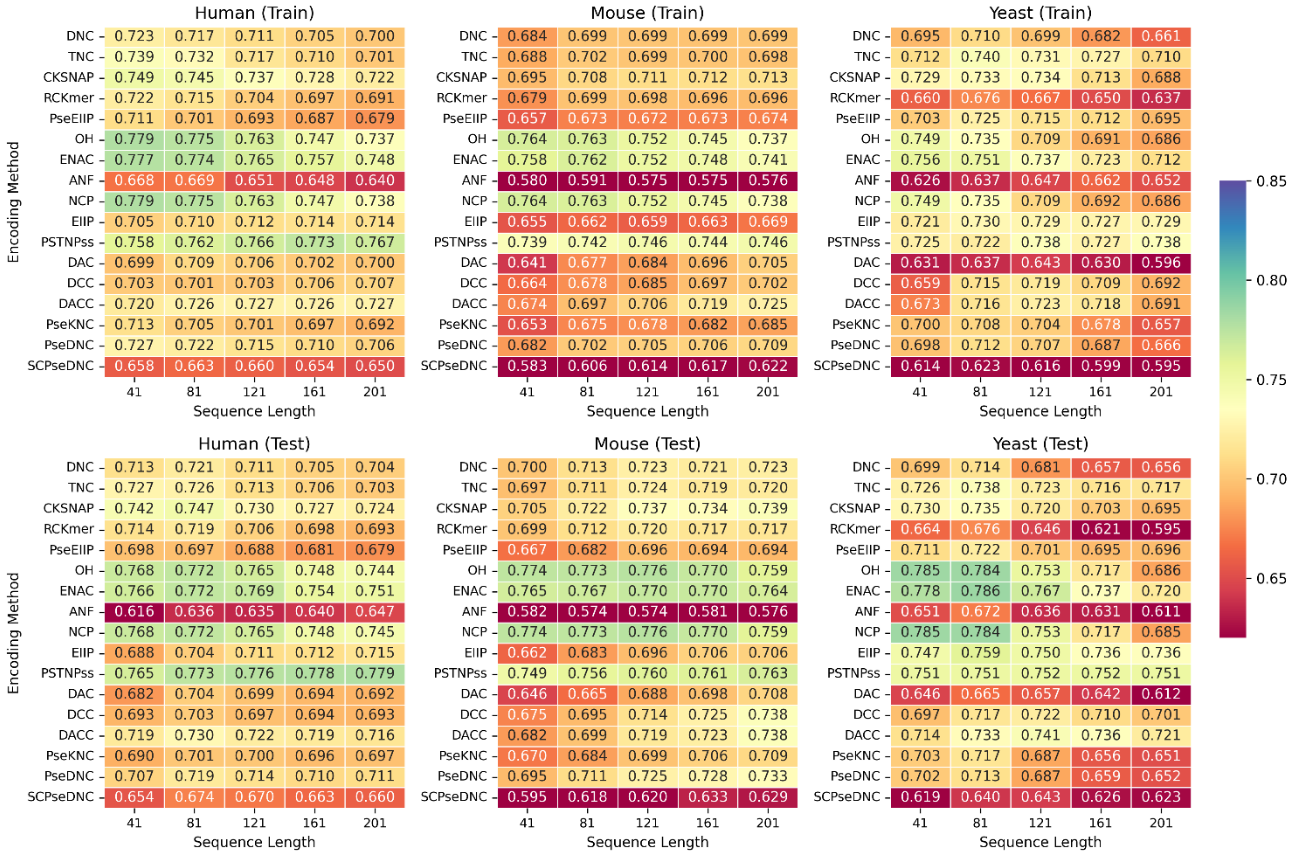


Figure S6 Comparative AUC performance of LR baseline models using 17 encoding methods across varying sequence lengths in human, mouse, and yeast.
